# Metamers of Bayesian computation

**DOI:** 10.1101/2020.08.11.246355

**Authors:** Hansem Sohn, Mehrdad Jazayeri

**Author notes:** **Correspondence** Mehrdad Jazayeri, Ph.D., Robert A. Swanson Career Development Professor, Assistant Professor, Department of Brain and Cognitive Sciences, Investigator, McGovern Institute for Brain Research, Investigator, Center for Sensorimotor Neural Engineering, MIT 46-6041, 43 Vassar Street, Cambridge, MA 02139, USA.

## Abstract

There are two sharply debated views on how humans make decisions under uncertainty. Bayesian decision theory posits that humans optimize their behavior by establishing and integrating internal models of past sensory experiences (priors) and decision outcomes (cost functions). An alternative model-free hypothesis posits that decisions are optimized through trial and error without explicit internal models for priors and cost functions. To distinguish between these possibilities, we introduce a novel paradigm that probes sensitivity of humans to transitions between prior-cost pairs that demand the same optimal policy (metamers) but distinct internal models. We demonstrate the utility of our approach in two experiments that were classically explained by model-based Bayesian theory. Our approach validates the model-based strategy in an interval timing task but not in a visuomotor rotation task. More generally, our work provides a domain-general approach for testing the circumstances under which humans implement model-based Bayesian computations.

## Introduction

Our daily life is filled with uncertain scenarios in which we have to evaluate different sources of information, infer the state of the world, and decide between different courses of action. For example, in the wake of COVID-19, we have to gather information about our surroundings to decide whether and when to eat out. This decision, like almost every other decision we make, depends on our prior beliefs (e.g., how probable it is to contract the virus), the present situation we face (e.g., how crowded the restaurant is), and the potential outcomes we expect (e.g., how bad it would be to get sick). Characterizing how humans represent and integrate these different sources of information is of utmost importance both for a deeper understanding of the nature of human cognition and for evaluating the conditions in which the decision process may go awry.

Bayesian decision theory (BDT) provides a normative framework for how to make optimal decisions under uncertainty^1–4^. Imagine an agent whose decision has to be adjusted based on the state of the environment. According to BDT, the optimal decision policy can be computed from three sources of information (Figure 1a): the prior probability distribution of the underlying state, the likelihood function of the state derived from noisy sensory evidence, and a cost (or reward) function that specifies possible decision outcomes. Optimal integration of these ingredients would enable the agent to maximize its expected reward (or minimize expected cost).

**Figure 1.**
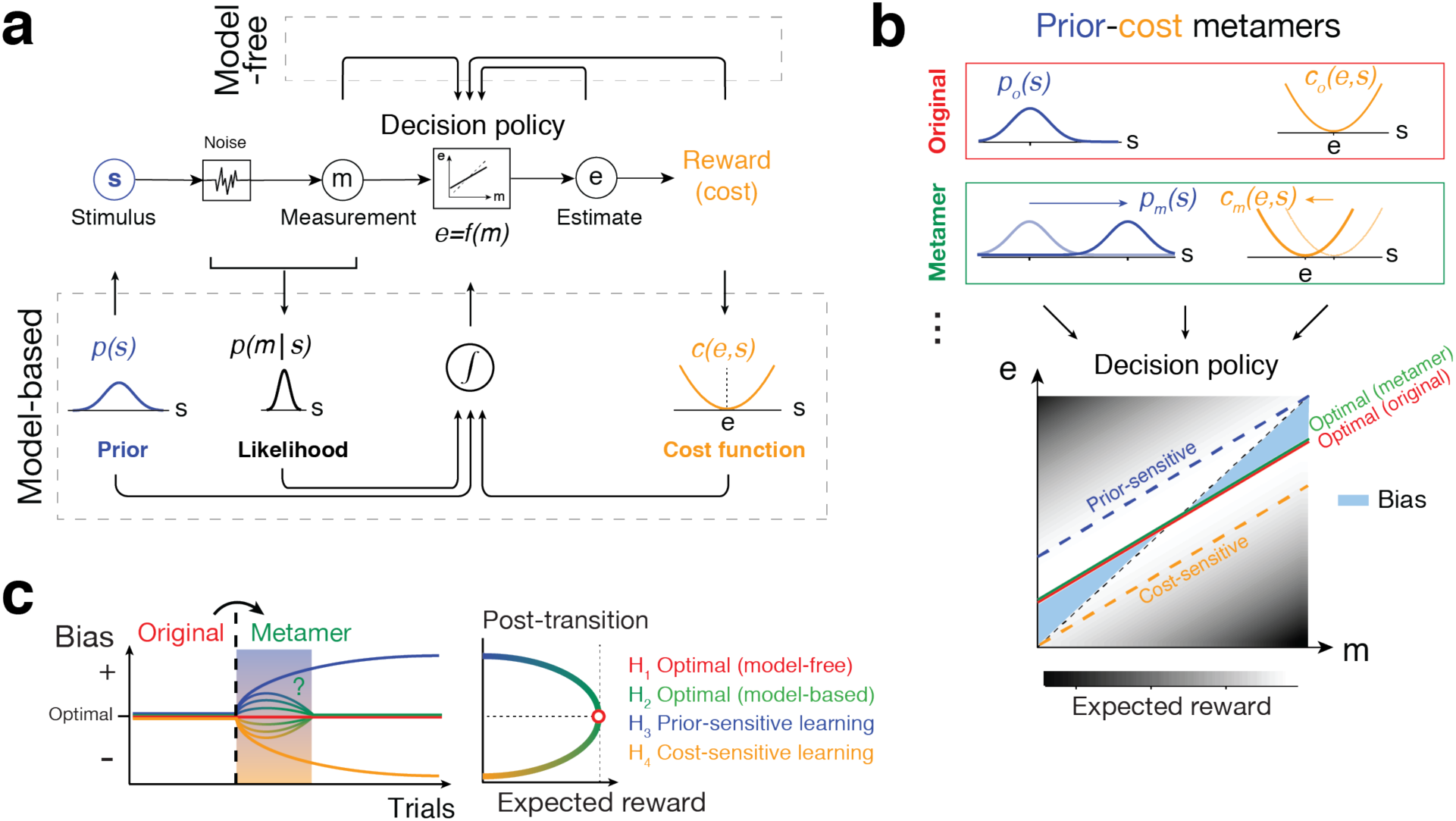
Testing Bayesian models of behavior using prior-cost metamers. **a**. Ideal observer model. An agent generates an optimal estimate (*e*) of stimulus (*s*) based on a noisy measurement (*m*). To do so, the agent must compute the optimal decision policy (rectangular box; *e=f*(*m*)) that maximizes expected reward. The decision policy can be computed using either a model-free (top) or model-based (bottom) learning strategy. In model-free learning, the policy is optimized through trial and error based on measurements and decision outcomes (arrows on top). In model-based learning, the agent derives the optimal policy by forming internal models for the stimulus prior probability, *p*(*s*), the likelihood of the stimulus after measurement, *p*(*m*|*s*), and the underlying cost function, *c*(*e,s*). **b**. Prior-cost metamers are different *p*(*s*) and *c*(*e,s*) pairs that lead to the same decision policy. Red box: An original pair (subscript *o*) showing a gaussian prior, *p*_*o*_*(s)*, a quadratic cost function, *c*_*o*_*(e,s)*. Green box: A metameric pair (subscript *m*) whose gaussian prior, *p*_*m*_*(s)*, and quadratic cost function, *c*_*m*_*(a,s)*, are suitably shifted to the right and left, respectively. Bottom: The optimal decision policy (*e=f*(*m*)) associated with both the original (red) and metamer (green) conditions is shown as a line whose slope is less than the unity line (black dashed line; unbiased). The colored dashed lines show suboptimal policies associated with an agent that is only sensitive to the change in prior (blue) or only sensitive to the change in the cost function (orange). The policy is overlaid on a gray-scale map that shows expected reward for various mappings of *m* to *e*. **c**. Hypothetical behavioral results after an uncued transition (vertical dashed line) from the original pair to its metamer. H_1_ (model free): After the transition, the agent continues to use the optimal policy associated with the original condition. Since this is also the optimal policy for the metamer, response biases do not change and expected reward remains at the maximum level (right). H_2_ (model-based): Immediately after the transition, the agent has to update its internal model for the new prior and cost function. This relearning phase causes a transient deviation from the optimal bias (curved lines after the switch, left) and reduces expected reward (right). After learning, the behavior becomes optimal since the optimal policy for the metaner is the same as the original. H_3_ (prior-sensitive): The agent only learns the new prior, which leads to a suboptimal behavior and reduced expected reward. H_4_ (cost-sensitive): Same as H_3_ for an agent that only learns the new cost function.

Remarkably, BDT can capture human behavior in a variety of domains including perception^5–10^, sensorimotor function^11–14^, multimodal integration^15–18^, and high-level cognitive function^19–21^. Based on the remarkable success of BDT, it has been proposed that the brain performs Bayesian computations by explicitly representing the sensory likelihood, prior knowledge, and cost function^22–26^.

However, the notion of the Bayesian brain is fiercely debated^27–29^. It is argued that the success of Bayesian models is unsurprising given the degrees of freedom researchers have in choosing the prior distribution, likelihood function, and cost function. The crux of the disagreement is about the value of formulating the optimization process in terms of a specific prior distribution and a specific cost function given that learning these components is not essential for learning the optimal policy^22,30,31^. A perfectly reasonable model-free alternative (Figure 1a) is that humans use trial-by-trial observations to incrementally arrive at the optimal solution without explicit reliance on the prior distribution and/or cost function^32–36^. Moreover, from a theoretical perspective, the choice of the prior and cost function is not unique. For example, an optimal agent may choose an option more frequently either because it is more probable or because it is more rewarding. More generally, in decision-making tasks, there are usually numerous pairs of priors and cost functions that, when combined, could lead to indistinguishable decision policies (Figure 1b). Analogous to the notion of metamers in perception^37–39^, we will refer to such pairs of priors and cost functions as prior-cost metamers. Because of the existence of such metamers, it remains an important and unresolved question whether decisions are made based on independently learned priors and cost functions.

Here we turn the problem of prior-cost metamers on its head to develop a general experimental strategy to test whether human decisions rely on independently learned priors and cost functions. The key idea behind our approach is to ask whether human behavior in a decision making task exhibits signs of relearning when we covertly switch from one prior-cost pair to another pair that is associated with exactly the same optimal decision policy (Figure 1c). According to BDT, optimal behavior depends on having learned the prior and cost function independently. Under this model-based hypothesis, changing to a new pair would lead to a transient change in decision policy until the observer relearns the new pair. Alternatively, if the optimal behavior were to emerge through trial-and-error without learning the prior and cost, then the observer should show no sensitivity to the switch and the behavior should remain optimal. We applied this approach to two tasks – a time-interval reproduction task^40,41^ and a visuomotor rotation task^42^ – both of which were classically explained in terms of BDT. Our results substantiated the role of independently learned priors and cost functions in the timing task, but not in the visuomotor rotation task. Accordingly, future behavioral studies can take advantage of our approach based on metamers to rule out the possibility of non-Bayesian strategies in Bayesian-looking behaviors.

## Results

Although we can use Newton’s Laws to explain how an apple falls from a tree, we would not conclude that the apple ‘implements’ Newton’s Laws. In the same vein, the fact that we can explain a person’s behavior in terms of the Bayesian decision theory (BDT) does not necessarily mean that their brain relies on explicit representations of priors and cost functions. Here, we apply a new experimental approach to investigate whether Bayesian computations rely on explicit knowledge about priors and cost functions. We explore this question in the context of two behavioral tasks, a time-interval reproduction task and a visuomotor rotation task. Previous work has shown that human behavior in both tasks is nearly optimal and can be explained in terms of integrating priors and cost functions^40,42^. Here, we test whether optimal performance in these tasks can indeed be attributed to explicit reliance on priors and cost functions.

Our experimental approach for both tasks is the same. We start the experiment with a specific choice of prior and cost function; after performance saturates, we covertly switch to a prior-cost metamer associated with the same optimal policy. The key question we ask is whether the transient behavior immediately after the switch reflects any sign of relearning the new prior-cost pair. We use the transient behavior to distinguish between two hypotheses. Under one hypothesis (H_1_, Figure 1c), the optimal behavior emerges through trial-and-error without explicit learning of the prior and cost. We will refer to this strategy as model-free learning since the optimality does not rely on an explicit model of the prior and/or cost. This hypothesis predicts that behavior shows no sensitivity to the switch and remains optimal because the optimal policy is the same as the one associated with the originally learned prior-cost pair. Under another hypothesis (H_2_), optimal behavior depends on learning the prior and cost as prescribed by the BDT. We will refer to this strategy as model-based learning since the agent has to build an internal model for the prior and cost. This hypothesis predicts that the behavior transiently deviates from optimality while the new prior and cost are being learned, but the asymptotic behavior would be optimal, as H_1_. For comparison, we will also test the two additional hypotheses, one in which the behavior is assumed to be solely sensitive to the prior (H_3_), and one in which the behavior is assumed to be solely sensitive to the cost function (H_4_). These hypotheses provide an informative benchmark to examine whether participants exhibit differential sensitivity to the prior or cost function.

### Time-interval reproduction task: Ready-Set-Go (RSG)

In the RSG task (Figure 2a), participants measure a time interval between the first two beats of an isochronous rhythm (‘Ready’ and ‘Set’ flashes) and are asked to press a button at the expected time of the third omitted beat (‘Go’). We refer to the sample interval between Ready and Set as *t*_*s*_, and the production interval between Set and Go as *t*_*p*_. We evaluated performance in RSG by what we refer to as a ‘metronomic function’, which quantifies *t*_*p*_ as a function of *t*_*s*_. On each trial, *t*_*s*_ is sampled from a prior probability distribution and participants receive graded numeric feedback as reward (*R*) depending on the absolute error between *t*_*p*_ and *t*_*s*_. We set the value of *R* according to a truncated quadratic function with a maximum of 100 and a minimum of 0 (Figure 2c). We will use the terms reward and cost function interchangeably to refer to the functional relationship between R and error (*t*_*p*_-*t*_*s*_).

**Figure 2.**
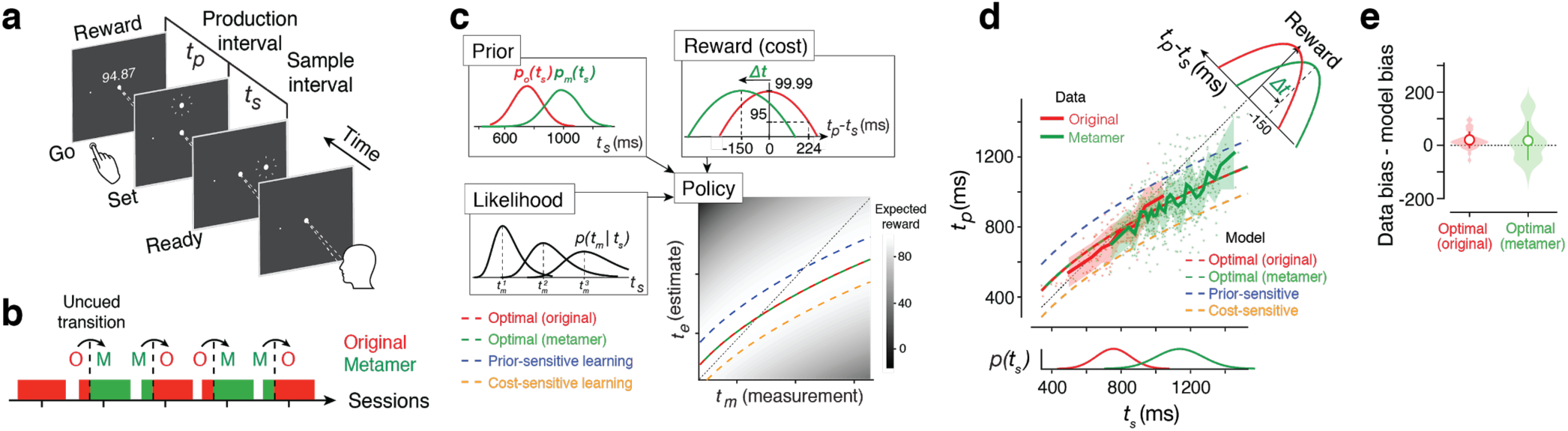
Prior-cost metamers in the Ready-Set-Go task. **a**. Time-interval reproduction task (“Ready-Set-Go”). While looking at a central spot, participants measure a sample interval (*t*_*s*_) demarcated by two flashes (Ready and Set), and initiate a delayed button press (Go) to produce a matching interval (*t*_*p*_) immediately after Set. At the end of each trial, participants received a numerical score whose value was determined by a cost/reward function (see below). **b**. Experimental sessions. In the first session that served as a baseline, participants performed the task with the original prior-cost set (‘O’). In the subsequent 4 sessions, we introduced an uncued transition between the original and metamer (‘M’) in an alternating fashion. In these sessions, the pre- and post-transition included ∼170, and ∼500 trials, respectively. In all panels we use red for the original condition and green for the metamer. **c**. Prior-cost metamers. Top-left: the original prior, *p*_*o*_(*t*_*s*_), is a Gaussian distribution (mean: 750 ms, SD: 144 ms) and its metamer, *p*_*m*_(*t*_*s*_), is a mixture of Gaussian distribution (see Methods). Top-right: reward (or cost) function is an inverted quadratic function of the error, *t*_*p*_*-t*_*s*_, that is truncated to have only positive values. The original cost function is centered at zero error, and the metamer is the same function shifted by *Δt*. Bottom-left: The likelihood function for *t*_*s*_ based on noisy measurement, *t*_*m*_, denoted *p*(*t*_*m*_|*t*_*s*_). The likelihood function is asymmetric and broader for longer *t*_*s*_ because measurement noise is assumed to scale with *t*_*s*_. Bottom-right: The optimal policy function that prescribes how an ideal observer should derive an estimate, *t*_*e*_, from *t*_*m*_. By design, the optimal policy for the original (H_1_) and metamer (H_2_) are identical. The plot also shows suboptimal policies for a prior-sensitive (blue; H_3_) and cost-sensitive (orange; H_4_) overlaid on a grayscale map showing average expected reward for different mapping of *t*_*m*_ to *t*_*e*_. When the original pair switches to its metamer, H_3_ and H_4_ predict positive and negative offsets, respectively, relative to the optimal policy. **d**. Data from a representative session showing *t*_*p*_ as a function of *t*_*s*_ (dots: individual trials, solid lines: running average across trials; shaded area: SEM). Prediction from H_1_ to H_4_ are shown in the same format as panel c. Below: The original and metamer priors. Top-right: The original and metamer cost functions. **e**. Bias in the original and metamer contexts. Violin plot showing the difference between the observed and optimal bias across all participants and all sessions with transitions (N=40, shaded region: distribution, open circles: grand averages; error bars: standard deviation). Bias was computed as the root mean squared difference between running average of *t*_*p*_ and *t*_*s*_.

Before analyzing the participants’ behavior, we characterized the predictions of various hypotheses regarding behavior in the context of two metameric pairs. Under both H_1_ and H_2_, the behavior is expected to be asymptotically optimal (i.e., after learning). We determined the form of the optimal policy by characterizing the behavior of an ideal observer performing the RSG task. An ideal observer integrates the likelihood function associated with the measured interval (*t*_*m*_) with the prior and the cost function to derive an optimal estimate (*t*_*e*_) that maximizes expected reward. This Bayes-optimal integration manifests as a nonlinear relationship between *t*_*e*_ and *t*_*m*_ (Figure 2c, bottom right). Accordingly, the *t*_*p*_ values for a participant that employs an optimal strategy should exhibit characteristic biases toward the mean of the prior. At first glance, the bias in the optimal policy may seem counter-intuitive. However, biasing responses in this way reduces the variance of responses such that the performance improvement due to reduced variance is larger than the performance drop due to the addition of biases. We also predicted the behavior for the prior-sensitive (H_3_) and cost-sensitive (H_4_) hypotheses after the switch to its metamer. According to H_3_, *t*_*p*_ values should exhibit an overall positive bias toward longer intervals. In contrast, under H_4_, *t*_*p*_ values should exhibit an overall negative bias. Moreover, since both hypotheses are suboptimal, they predict an overall drop in expected reward.

Next, we collected data from 11 participants performing the RSG task in the context of two distinct metameric prior-cost pairs across 5 daily sessions (Figure 2b). In the first session, participants performed the task under the ‘original’ pair, which consisted of a Gaussian prior distribution and a cost function that was centered at zero when *t*_*p*_ = *t*_*s*_ (Figure 2c; red). For the original pair, participants’ *t*_*p*_ increased with *t*_*s*_, and exhibited systematic biases toward the mean of the prior (Figure 2d, red). Similar to numerous previous studies^40,41,43–46^, this behavior was consistent with predictions of the ideal observer model (Supplementary Figure 1).

Our goal for the subsequent four sessions was to test participants’ behavior after a covert switch between the original pair and its ‘metamer’. To design the metamer (green in Figure 2c), we used the same truncated quadratic form for the cost function but shifted it by −150ms (*Δt*) such that the maximum value of *R* was now associated with responding 150 ms earlier than the third beat (*t*_*p*_ = *t*_*s*_ - 150ms). Next, we designed the metameric prior. Intuitively, the new prior has to be shifted in the positive direction to counter the negative shift in the cost function and lead to the same decision policy. However, to achieve a perfect metamer, the new prior cannot be Gaussian (see Methods). We modeled the new prior as a Gaussian Mixture Model (GMM) whose parameters were adjusted such that the integration of the GMM and shifted cost function produced the same optimal policy. Note that the exact form of the GMM depends on measurement noise level and was therefore customized for each participant independently (see Methods). For each subject, we verified that the optimal policy for the original and metamer were indeed the same (Supplementary Figure 2).

Participants’ behavior in the context of the metamer showed similar biases toward the mean (Figure 2d, green; see Supplementary Figure 2 for all subjects) and was asymptotically matched to that of the ideal observer (Figure 2e, green; p=0.137, t test for equal bias between data and optimal model). In other words, participants’ behavior during both the original and its metamer was captured by the same optimal policy. This finding is consistent with both the model-free (H_1_) and model-based (H_2_) optimal strategy.

Our main interest, however, is to distinguish between the model-free (H_1_) and model-based (H_2_) strategies. Recall that any transient deviation from the behavior of an ideal observer after the switch would provide evidence against the model-free strategy of using the original policy (no re-learning). Therefore, we estimated the log-probability of behavioral data under the ideal-observer model before, immediately after, and long after the covert switch. As expected, the magnitude of bias across the participants was nearly matched to that of an ideal observer before the switch (Figure 3a, right inset). After the transition, the probability of data under the ideal observer model dropped sharply and transiently (Figure 3a, left), and within approximately 50 trials, moved back to the pre-transition level (nonparametric Friedman test for effect of transition on mean p(optimal policy) across [-25 0], [1 25], [26 50] trials with respect to the transition, p=0.020; post-hoc signed rank test for 25 trials before versus after the transition, p=0.081; post-hoc signed rank test for 1-25 trials versus 26-50 trials after the transition, p=0.003). It is possible that the transient deviation from optimality was due to the fact that participants experienced new *t*_*s*_ values after the prior was switched. We ruled out this possibility by analyzing only the trials whose *t*_*s*_ values overlapped with the distribution of *t*_*s*_ before the transition (Supplementary Figure 3). Note that the transient deviation from the ideal observer behavior was small in comparison to what is expected from a purely prior-sensitive (H_3_; Figure 3b) or cost-sensitive (H_4_; Figure 3c) strategy. Together, these results reject H_1_, H_3_ and H_4_, but not H_2_, suggesting that participants were indeed sensitive to both the prior and cost function despite the metameric relationship between the prior-cost pairs before and after the switch.

**Figure 3.**
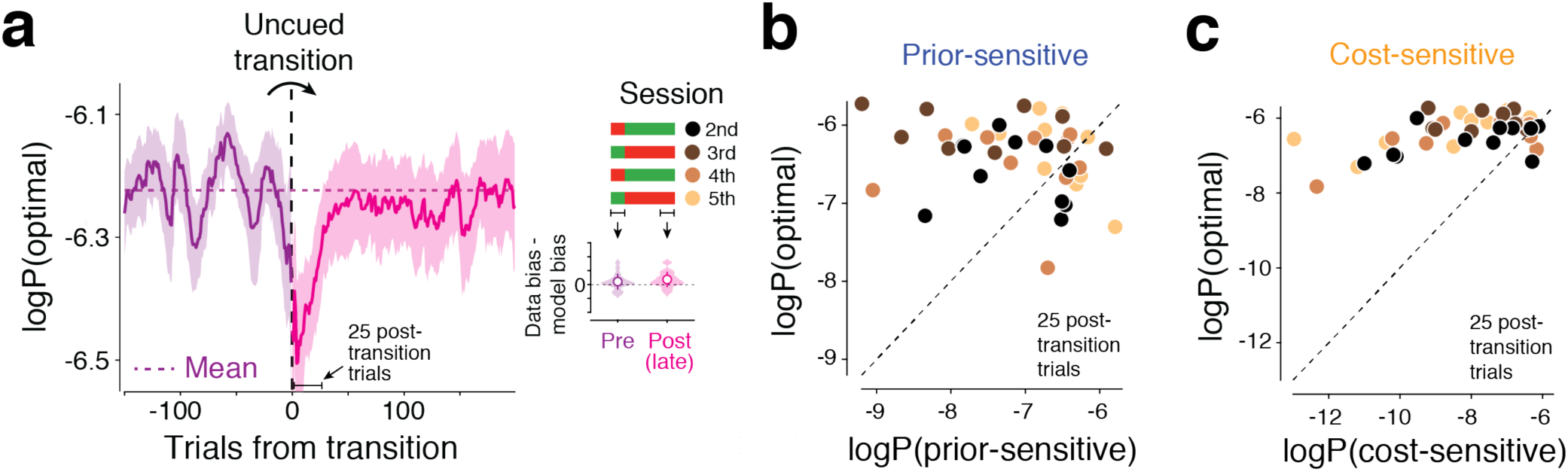
Behavior immediately after the switch to a new prior-cost condition in the RSG task. **a**. Log probability of the optimal policy, denoted ‘logP(optimal)’ under the data across a few hundred trials before and after the transitions (vertical dashed line) averaged across subjects and sessions (N=40, shaded area: SEM, purple: pre-transition, magenta: post-transition; horizontal dotted line: log probability of the optimal policy averaged across 100 pre-transition trials). Data from individual trials are smoothed using a causal Gaussian kernel (SD: 10 trials), separately for the pre- and post-transition. Twenty five post-transition trials are also highlighted as a window of interest for later analyses. Legend illustrates the alternating transitions between the original (red) and metamer (green) across 4 sessions. Right inset: the average difference between observed and optimal bias in the pre- and post-transition epochs plotted in the same format as Figure 2e. Pre-transition includes all trials before transition. Post-transition includes the last 200 trials (to avoid misestimation due to the transient). **b**. Comparison of the optimal policy, logP(optimal), with the suboptimal prior-sensitive policy, denoted logP(prior-sensitive), based on the data in the 25 post-transition trials across participants and sessions (N=40; see legend in **a**). **c**. Same as **b** for the cost-sensitive model.

### Learning dynamics of the prior-cost metamer

So far, we have established that participants were sensitive to transitions between prior-cost metamers, which suggests an underlying dynamic process for learning the new prior and cost function. To characterize this dynamic process, we sought to track the participants’ internal estimate of the prior and cost function during the learning process. To do so, we built an observer model in which the prior and cost function could change dynamically during learning. From a statistical (fitting) perspective, parameterizing the full prior distribution and cost function throughout learning is not feasible. We therefore built a simplified model in which the profile of the prior and cost function were the same as those associated with the original pair (i.e., Gaussian and quadratic, respectively). However, we allowed the mean (*µ*) and standard deviation of the prior, and the shift in the cost function (*Δt*) in the model to be determined dynamically based on the observed behavior. We will use ‘subjective prior’ and ‘subjective cost function’ to refer to *µ* and *Δt*, respectively.

We fitted this observer model to each participant’s behavior using a running window of 100 trials and tracked the fits to subjective prior and cost function from before to after the transition. Across participants, *µ* and *Δt* changed systematically and in accordance with the changes in the experimentally imposed prior and cost function (Figure 4a,b; see Supplementary Figure 4 for the standard deviation of the prior). We also use Bayesian Information Criterion (BIC) to evaluate the data with respect to the model-free (H_1_), model-based (H_2_), prior-sensitive (H_3_), and cost-sensitive (H_4_) hypotheses (Figure 4c). Results provided clear evidence in support of the model-based hypothesis. Together, these analyses substantiate the presence of a dynamic learning process for the prior and cost function after the transition, and provides an estimate of the underlying time course of learning in this task.

**Figure 4.**
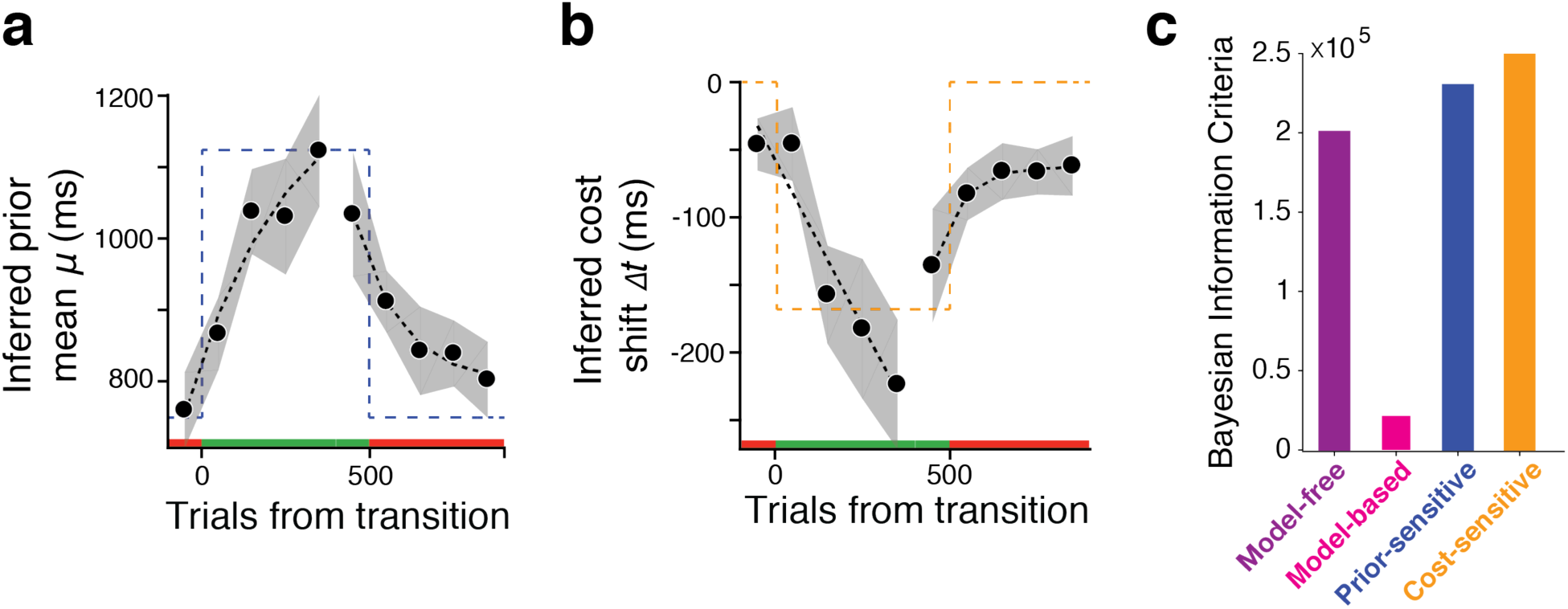
Time course of learning the new prior and cost function across subjects. **a**,**b**. The inferred mean of the prior (*µ*) during two types of transitions between the original and metamer conditions (red and green above the abscissa). The mean was inferred from fits of an observer model to behavior in running windows of 100 trials at different time points before and after the transition (black circles: average across participants and sessions N=40, shades: SEM, black dotted line: exponential fit; dashed line: the experimentally imposed prior mean). **b**. The inferred shift in the subjective cost function (*Δt*) during transitions between the original and metamer conditions shown with the same format as **a**. **c**. Model comparison using Bayesian information criteria (BIC). Smaller BIC values indicate the better models.

One important observation from the learning dynamics was that the prior and cost function changed in parallel and their learning rates appeared to be comparable (Figure 4a,b). Indeed, fitting the learning curve for the prior and cost function with an exponential function led to comparable learning rates (Figure 5a; p=0.42, signed rank test). To gain a deeper understanding of the computational consequences of this parallel learning we performed a series of simulations in which we varied the learning rate for the prior and cost function (Figure 5f). The simulations indicated that when the prior learning was faster than the cost function, the behavior after the transition became more like the prior-sensitive model (H_3_) and led to a reduction of expected reward. A higher learning rate for the cost function also reduced performance by causing the behavior to become more like the suboptimal cost-sensitive model (H_4_). The best performance during transition was associated with cases when two learning rates were comparable. This result provides a normative argument that it may be beneficial to have comparable learning rates for the prior and cost.

**Figure 5.**
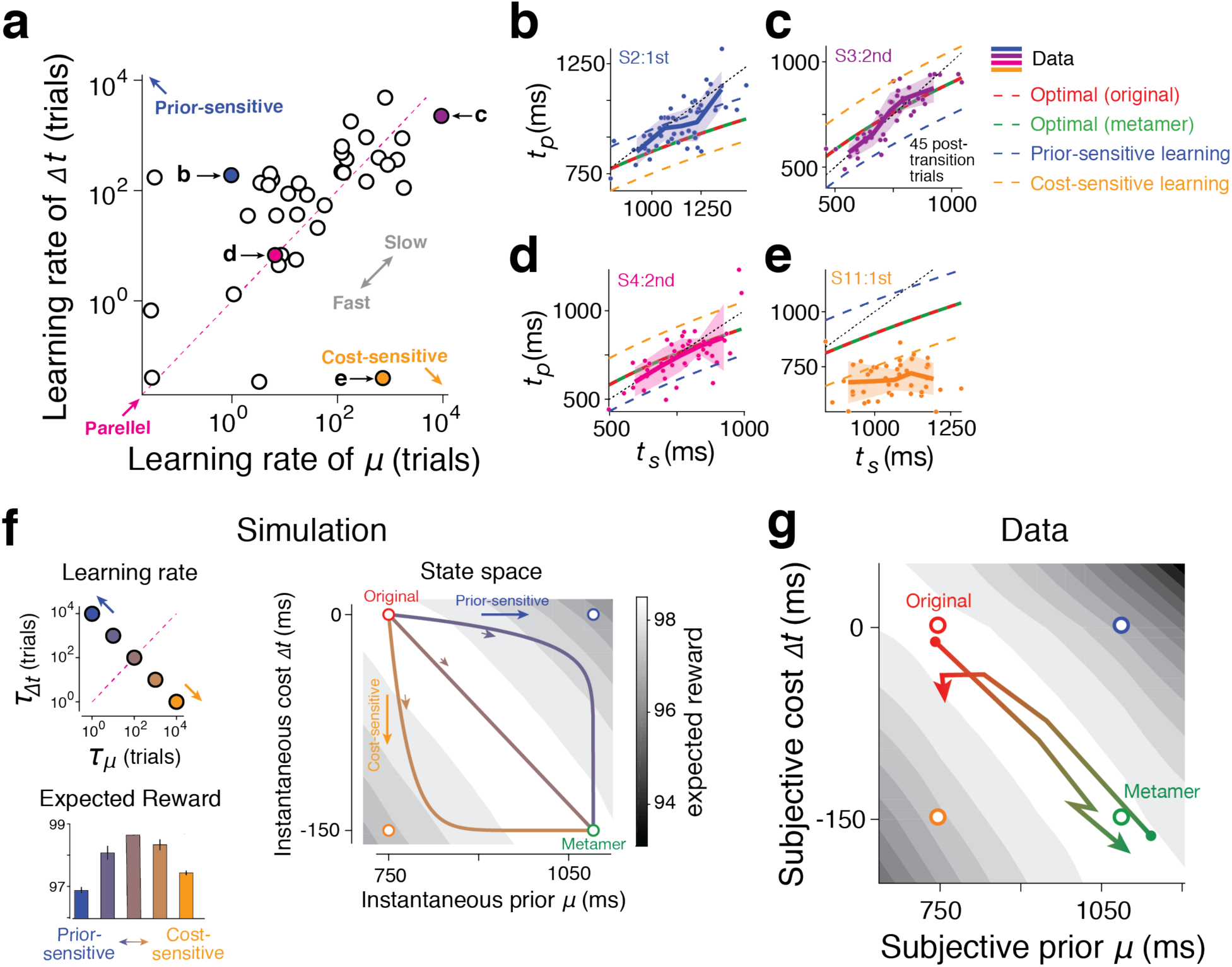
Parallel learning of prior and cost function and its computational consequences. **a**. Learning rate for adjusting the mean of the prior (*µ*) and the shift in the cost function (*Δt*) after prior-cost transitions. We inferred the learning rates by fitting exponential curves to learning dynamics (Figure 4). For many participants, the learning rates were comparable as evident by the points near the unity line (‘parallel’, magenta dotted line). Points above the unity line indicate faster adjustment for *µ* relative to *Δt*, which is expected from a participant that is more prior-sensitive (blue circle). Points below the unity line indicate faster adjustment for *Δt* relative to *µ*, which is expected from a participant that is more cost-sensitive (orange circle). Two outlier data points on top-right were not included in the plot (N=40 across subjects and sessions). **b-e**. Behavior of four example participants in the first 45 post-transition trials (dots: individual trials; solid line: average across bins; shaded region: SEM). The title of each plot indicates the participant (e.g., S2) as well as the session with transitions (e.g. ‘2nd’), and the colors correspond to the specific points in panel **a**. For each example, the predictions from various optimal and suboptimal models are also shown (legend). **f**. Simulation with varying learning rates. Top-left: We selected 5 pairs of the learning rates for *µ* and *Δt*, indexed by the corresponding time constants, *τ*_*µ*_, and *τ*_*Δt*_, respectively. The pairs ranged from faster prior learning (blue) to parallel learning (brown) to faster cost learning (orange). Right: Learning trajectory of *µ* and *Δt* for three example points in the top-left panel (with the corresponding colors) during the transition from the original prior-cost set (red circle) to its metamer (green circle). Faster prior learning first moves the state toward the prior-sensitive model (open blue circle) and then toward the end point. Faster cost learning first moves the state toward the cost-sensitive model (open orange circle). When the learning rates are identical (brown), the state moves directly from the original to the metamer. The grayscale colormap shows expected reward as a function of *µ* and *Δt* (Equation 19 in Methods). Bottom-left: Average reward along the learning trajectory for the points in the top-left panel. Results correspond to the averages of 500 simulated trials (error bar: SEM). **g**. Learning trajectories across participants based on average learning rates in panel **a** shown separately while transitioning from original to metamer (red to green), and from metamer to original (green to red). Note that, on average, the expected reward (grayscale colormap, same as panel (f)) remains high.

To address the question, we performed a simulation to estimate the expected reward in a state space (Figure 5f,g) comprised of the mean of the subjective prior (*µ*) and the shift in the cost function (*Δt*). As expected, when the subjective prior mean and cost shift match the objective values (original and metamer in Figure 5f,g), the expected reward is maximum. Any other point in the space would give rise to suboptimal reward amounts. However, we unexpectedly found a continuum of the prior-cost pairs that can achieve almost the maximum reward, in between the original and metamer sets we devised. Therefore, if the subjects learned the new prior and cost in parallel – i.e., navigating the state space diagonally, the expected reward would not decrease as subjects remain in the continuous regime of the optimal reward. However, the reward would decline rapidly as subjects sequentially update their internal prior or cost function.

One intriguing conclusion from these simulations is that when prior and cost learning are suitably coordinated, the behavior may remain optimal throughout the relearning process. In other words, a participant’s behavior may show no sign of learning (i.e., no deviation from ideal observer model) even while they are in the process of learning the new prior and cost function. With this consideration in mind, we analyzed the behavior of individual participants asking whether such coordinated learning for the prior and cost function was evident in their behavior (Figure 5b-e). Results revealed a diversity of learning rates for the prior and cost function ranging from relatively faster prior learning (Figure 5b), to comparable learning rates for the prior and cost function (Figure 5c,d), to faster cost learning (Figure 5e). However, there was no systematic difference between the two learning rates across participants (Figure 5a).

The diversity of learning rates for the prior and cost function across participants provides a coherent explanation for various findings that we originally found puzzling. First, it explains why we were able to infer the prior mean and the shift in the cost function across participants (Figure 4a,b). If the learning dynamics were identical across participants, we would have not been able to infer those dynamics because of the metameric relationship between the two conditions. Second, since the prior and cost learning rates were comparable across participants, the deviation from the ideal observer policy after prior-cost switches was small (Figure 3a) compared to the prior-sensitive and cost-sensitive strategies (Figure 3b,c). Third, the parallel learning enables participants to maintain a steady performance after prior-cost switches (Supplementary Figure 5).

### Visuomotor rotation (VMR) task

In the VMR task (Figure 6a), participants use a manipulandum to move a cursor from the center of a visible ring to the remembered position of a target flashed briefly on the circumference of that ring. As soon as the cursor begins to move, we make the cursor invisible and change its angular position relative to the angular position of the hand by a rotation angle, *x*_*s*_. While moving, the cursor reappears briefly when it is midway, which provides information about *x*_*s*_ to the participants. On each trial, *x*_*s*_ was sampled from a Gaussian prior distribution (Figure 6c), and a numerical feedback, *R*, was provided depending on the error between *x*_*s*_ and the corresponding correction, *x*_*p*_, which we defined as the angle between the hand position and the target on the ring. We set the value of *R* according to a truncated quadratic function with a maximum of 100 and a minimum of 0 (Figure 6c).

**Figure 6.**
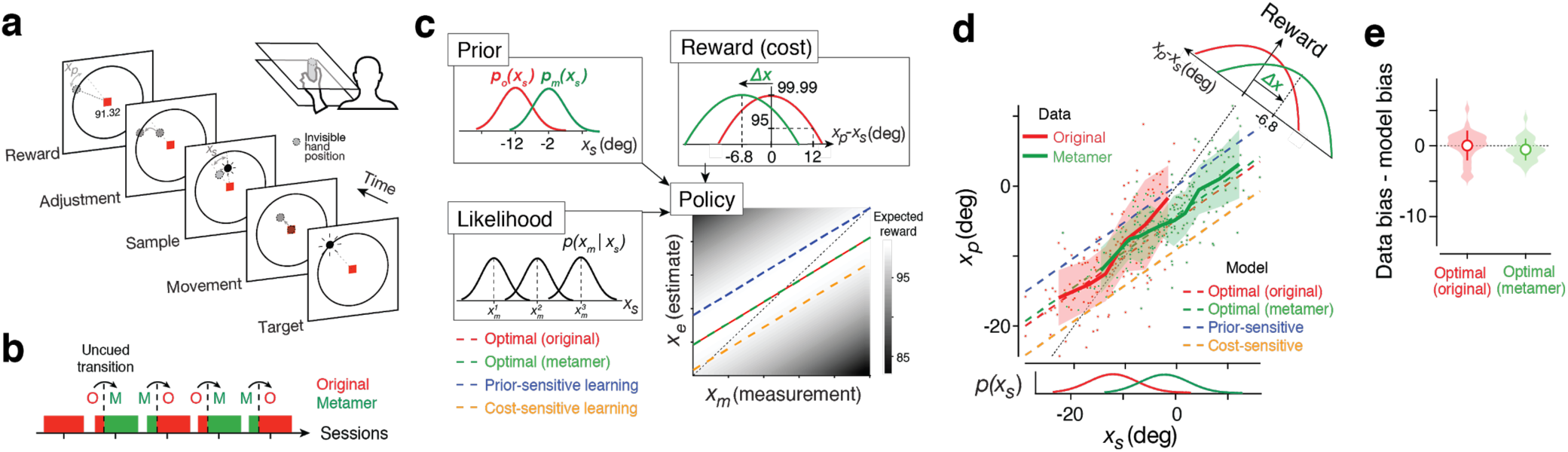
Prior-cost metamers in the visuomotor rotation task. **a**. Visuomotor rotation task. Participants move a manipulandum (vertical cylinder) to maneuver a cursor on a horizontal display (top-right) from the center of the screen (red square) to a target flashed briefly on the circumference of a visible ring (‘Target’: black circle on the ring). The moving cursor is occluded while moving except from a brief reappearance halfway along the path (‘Sample’: black circle). The cursor position is covertly rotated by an angle (*x*_*s*_) relative to the position of the hand (gray circle). Participants have to use *x*_*s*_ and apply a matching counterrotation (*x*_*p*_) to bring the cursor to the target position (‘Adjustment’). At the end of each trial, participants received a numerical score whose value was determined by a cost/reward function (see below). **b**. Experimental sessions shown in the same format as Figure 2b. **c**. Prior-cost metamers shown in the same format as in Figure 2c. Top-left: the original prior, *p*_*o*_(*x*_*s*_), is a Gaussian distribution (mean: −12 deg, SD: 7.5 deg) and its metamer, *p*_*m*_(*x*_*s*_), is also a Gaussian distribution shifted by 10 deg (see Methods). Top-right: reward (or cost) function is an inverted quadratic function of the error, *x*_*p*_*-x*_*s*_, that is truncated to have only positive values. The original cost function is centered at zero error, and the metamer is the same function shifted by *Δt*. Bottom-left: The likelihood function for *x*_*s*_ based on noisy measurement, *x*_*m*_, denoted *p*(*x*_*m*_|*x*_*s*_). Bottom-right: The optimal policy function that prescribes how an ideal observer should derive an estimate, *x*_*e*_, from *x*_*m*_. By design, the optimal policy for the original (H_1_) and metamer (H_2_) are identical. The plot also shows suboptimal policies for a prior-sensitive (blue; H_3_) and cost-sensitive (orange; H_4_) overlaid on a grayscale map showing average expected reward for different mapping of *x*_*m*_ to *x*_*e*_. H_3_ and H_4_ predict positive and negative offsets, respectively, relative to the optimal policy. **d**. Data from a representative session showing *x*_*p*_ as a function of *x*_*s*_ shown in the same format as in Figure 2d. **e**. Bias in the original and metamer contexts (N=36) in the same format as in Figure 2e.

Similar to the RSG task, we characterized the predictions of various hypotheses regarding behavior in the context of two metameric pairs. Under the model-free (H_1_) and model-based (H_2_) Bayesian strategies, the behavior is expected to be asymptotically optimal (i.e., after learning). We determined the form of the optimal policy by characterizing the behavior of an ideal observer that integrates the likelihood function based on a noisy measurement, *x*_*m*_ with the prior and the cost function to derive an optimal estimate, *x*_*e*_, that maximizes expected reward (Figure 6c). With a Gaussian likelihood function, a Gaussian prior and a quadratic cost function, the optimal policy is to map *x*_*m*_ to *x*_*e*_ linearly with a bias toward the mean (i.e., slope of less than 1; see Methods). Under the prior-sensitive (H_3_) and cost-sensitive (H_4_) hypotheses, *x*_*p*_ values would exhibit an overall positive and negative bias, respectively (Figure 6c), and would lead to lower reward than expected under the optimal policy. Note that linearity of the optimal policy in the VMR task is due to the relatively simple form of the likelihood function (Gaussian with a fixed standard deviation) compared to the RSG task in which variability scales with the base interval.

We collected data from 10 of the participants who also performed the RSG task. Our experimental procedure was the same as in RSG: an original prior-cost pair in the first session, and alternations between the original and its metamer in the subsequent 4 sessions (Figure 6b). The original prior was centered at −12 deg, and the original cost function was centered at zero. Participants’ behavior exhibited biases toward the mean as expected by the optimal policy (Figure 6d for representative data; see Supplementary Figure 6 for all participants; Figure 6e, red, p=0.561, t test, Null: equal bias between data and optimal model).

Next, we designed the prior-cost metamer (Figure 6c). For the prior, we shifted the Gaussian distribution by 10 deg so that the new mean was at −2 deg. For the cost function, we derived the appropriate shift in the cost function (*Δt*) needed to create a metameric pair (i.e., matching optimal policy). Note that the shift in the cost function had to be customized separately for each participant depending on the measurement noise fits in the first session. Participants’ behavior in the context of the metamer showed similar biases toward the mean (Figure 6d, green; Supplementary Figure 6 for all participants) and was asymptotically matched to that of the ideal observer (Figure 2e, green; p=0.052, t test for equal bias for data and optimal model). In other words, participants used the same optimal policy for both the original and its metamer, consistent with both the model-free (H_1_) and model-based (H_2_) strategy.

To distinguish between the model-free (H_1_) and model-based (H_2_) strategy, we tested whether there was any transient change in decision policy immediately after the switch between the two prior-cost pairs. Unlike RSG, participants’ behavior did not exhibit any transient deviation from the optimal policy after the switch (Figure 7a, Supplementary Figure 7), and was better explained by the optimal policy than a purely prior-sensitive (H_3_) or cost-sensitive (H_4_) strategy, even with the first 25 trials after the switch (Figure 7b,c; Supplementary Figure 7). Together, these results suggest that, in VMR, participants rely on a model-free optimal policy (H_1_) without explicit learning of the prior and cost function. More generally, this results invites caution against interpreting Bayes-optimal behavior as evidence for knowledge about the underlying priors and the cost functions.

**Figure 7.**
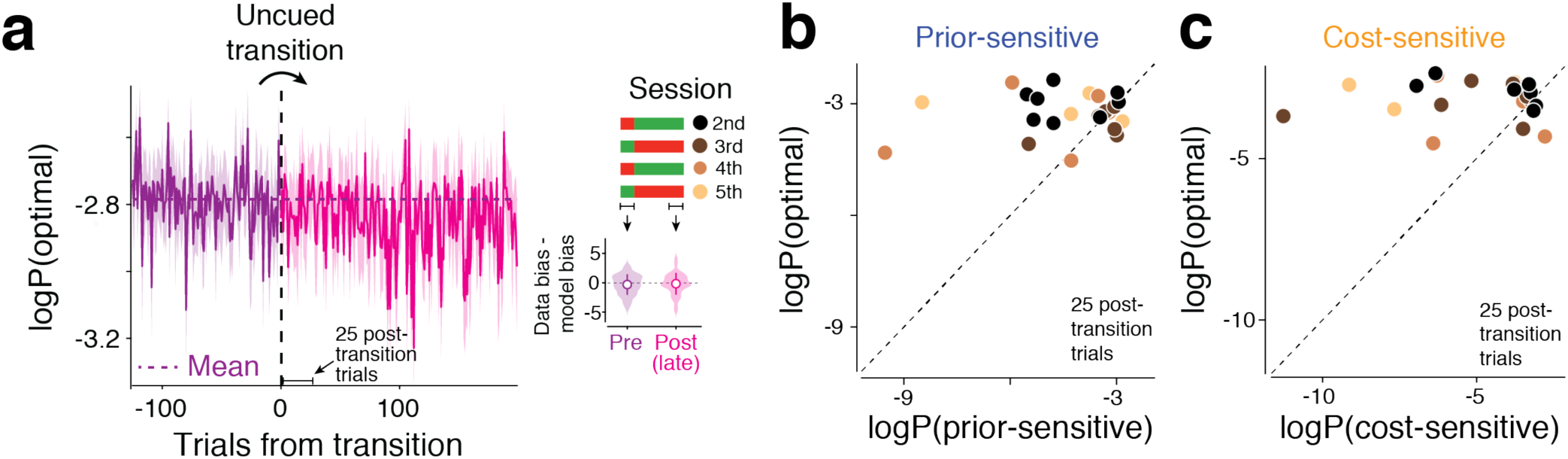
Behavior immediately after the switch to a new prior-cost condition in the VMR task. Results are shown in the same format as in Figure 3. Unlike the RSG task, behavior does not exhibit any transient deviation from optimal policy after the switch.

## Discussion

There is a strong consensus that human decision making under uncertainty is consistent with Bayesian decision theory (BDT)^1,3,4,22,25^. This finding has been taken as evidence that humans have internal models for the prior probability of environmental states, the likelihood of those states based on sensory evidence, and the costs (or rewards) associated with making decisions about those states. However, the success of fitting Bayesian models to behavior does not necessarily mean that the human brain establishes internal models for priors, likelihoods, and cost functions. Indeed, numerous researchers have contested the formulation of optimal behavior in terms of model-based BDT, arguing instead that the brain relies on heuristics^27,47^ and gradual learning of model-free optimal policies^29,35,36^.

There have been numerous attempts to experimentally test the key model-based prediction of BDT. For example, some studies have changed one element of the BDT during experiments and studied how human observers adapt to the change^41,48–50^. However, this approach inevitably changes the corresponding optimal decision policy and therefore could not be used to distinguish between model-based and model-free hypotheses. One potential approach for tackling this problem is to ask whether knowledge about the prior and cost would transfer to a new behavioral setting^32,33^. For example, one can expose participants to two prior-cost conditions, P_1_,C_1_, and P_2_,C_2_, and ask whether behavior remains optimal when participants are tested in cross conditions; i.e., P_1_,C_2_ and P_2_,C_1_. If task performance stays at the optimal level under the new pair, it indicates that participants can flexibly integrate learned priors and cost functions in accordance with BDT. However, implementation of this approach can be challenging since it requires participants to fully learn and flexibly access distinct priors and cost functions. As such, deviations from optimality using this approach may be due to bounded computational capacity that humans have in switching between internal models.

To fill this gap, we developed a novel experimental paradigm that capitalizes on the issue of model identifiability in BDT^51^. Since there is no one-to-one correspondence between an optimal decision policy and its underlying BDT elements (likelihood, prior, and cost function), it is possible to design Bayesian metamers that involve different prior-cost pairs but are associated with the same optimal policy. This approach parallels fruitful uses of metamers in perception^37–39^ and machine learning^52^. The key idea is that an observer whose decisions rely on a model-free optimal policy should ‘see’ the pairs as metamers because they correspond to the same policy. In contrast, an observer that makes direct use of priors and cost functions will see the two pairs as distinct and will therefore be sensitive to transitions between them.

We demonstrated the utility of our approach in the RSG and VMR tasks that humans perform nearly optimally. Switching between metamers led to transient deviations from the optimal policy in the RSG but not VMR task. These findings are consistent with the interpretation that the optimal policy in the RSG – not VMR – task relies on internal models for the prior and cost function. What factors might underlie the difference between the two tasks? One factor that differs between the two tasks is the complexity of the decision policies. Recall that the optimal policy in the VMR task was a linear mapping between measurements and estimates. It is conceivable that the sensorimotor system has powerful machinery to rapidly adapt to new linear sensorimotor mappings without the need to relearn internal models of the prior and cost function. In contrast, the nonlinear optimal policy in the RSG task may necessitate the re-learning of the prior and cost function for making adjustments to the policy. It may also be that the difference is due to inherent differences between sensorimotor timing (RSG) and sensorimotor reaching (VMR). More experiments involving different forms of priors, cost functions, and optimal policies are needed to understand the conditions under which the brain adopts model-based versus model-free strategies^53,54^.

An additional insight from our study of learning under metamers was the importance of learning dynamics. We found that when the learning rates for the prior and cost function are comparable, observers can learn new internal models without sacrificing performance during the learning. As a corollary, parallel learning may cause model-based learning to appear as model-free. Indeed, this observation reveals the Achilles’ heel of our paradigm: one would not be able to reject the model-based approach if participants’ learning rates for priors and cost functions are comparable. Therefore, our paradigm provides sufficient – not necessary – evidence for the model-based strategy.

Our framework can be flexibly extended in different ways. For example, one can further validate BDT using likelihood-cost, likelihood-prior metamers. The use of likelihood-cost metamers may be particularly valuable when the likelihood and cost can be manipulated^6,15,42^ but the prior is thought to be hard-wired, perhaps as a result of development and/or evolution^10^. For the prior-cost metamers that we focused on, there is a great deal of flexibility in choosing the parametric form of the prior and cost function. This flexibility can be used to validate conclusions that previous studies made about the role of a specific form of prior distribution^55^ and/or cost function^40,56^ in human sensorimotor behavior. Finally, the specific design considerations for the metamer is not limited to linear shifts of the prior and/or cost function and can be adjusted based on experimental needs.

Care must be taken in interpreting the results of metamertic experiments such as ours. For example, one implicit assumption of our methodology is that the model-free strategies do not involve updating the decision policy. This is a reasonable assumption if stimulus statistics associated with the original and metamer priors are sufficiently similar. However, if the stimuli associated with the new prior are widely different from the original prior, participants may have to engage in a learning process to establish the optimal decision policy for the new stimuli. This learning may present itself as a transient deviation from optimality and can thus be misinterpreted for model-based learning. This concern is unlikely to have impacted our interpretation since we made the shifts in the prior and cost function relatively small. Moreover, we found that participants’ behavior deviated from optimality even when we focused our analysis only on those stimulus conditions that were present in both the original and metalmer conditions (Supplementary Figure 3). However, not addressing this issue could lead to a misinterpretation of a model-free policy as model-based BDT. One strategy that can further reduce the chance of such misinterpretation is to design metamers that involve fully overlapping ranges of stimuli (i.e., have the same domain) but with different probability profiles (e.g., different variance or skewness). This strategy would ensure that participants have observed all possible stimuli in the metamer before the transition.

It is also important to carefully titrate the cost function. If the transition to the metamer leads to an overall increase or decrease in reward, participants might notice the change and initiate exploratory behavior to update the decision policy. Again, this concern is unlikely to have impacted our results since we adjusted the metamer cost function to ensure the overall expected reward was unchanged after the transition. Nevertheless, it is important to design the cost function metamers carefully so as to avoid misattributing learning of a new model-free policy to model-based learning.

Our analysis of learning dynamics in RSG suggests that updating priors and cost functions likely involve different neural systems. This conclusion is consistent with the underlying neurobiology^57^. Humans and animals update their prior beliefs when observed stimuli deviate from predictions, which can be quantified in terms of a sensory prediction error (SPE)^22,58–60^. The cerebellum is thought to play a particularly important role in supervised SPE-dependent learning of stimulus statistics in sensorimotor behaviors^61–64^. In contrast, learning cost functions is thought to depend on computing reward prediction errors (RPE) between actual and expected reward^65–67^. This type of learning is thought to involve the midbrain dopaminergic system ^68^ in conjunction with the cortico-basal ganglia circuits^57,69–71^. Finally, the information about the prior and cost function has to be integrated to drive optimal behavior. Currently, the neural circuits and mechanisms that are responsible for this integration are not well understood^72–74^. Future work could take advantage of our methodology to systematically probe behavioral settings that rely on model-based BDT as a rational starting point for making inquiries about the underlying neural mechanisms.

## Methods

Eleven participants (age: 18-65 years, 6 male and 5 female) participated in the experiments after giving informed consent. All participants were naive to the purpose of the study, had normal or corrected-to-normal vision, and were paid for their participation. All experiments were approved by the Committee on the Use of Humans as Experimental Subjects at the Massachusetts Institute of Technology.

### Procedures

All 11 participants completed five experimental sessions of the Ready-Set-Go time interval reproduction task (RSG), and 10 of them completed five sessions of the visuomotor rotation task (VMR). The testing sequence for the two tasks was counterbalanced across participants. In each session, a participant was seated in a dark quiet room and asked to perform the task of interest for approximately 60 min. For both tasks, stimuli and behavioral contingencies were controlled by an open-source software (MWorks; http://mworks-project.org/) running on an Apple Macintosh platform.

Before the first session, participants received detailed instruction about task contingencies, and completed dozens of practice trials. We used the data from the first session to make baseline measurements of various participant-specific parameters needed for adjusting experimental parameters in the remaining sessions (see below for details). The data from the remaining sessions were used to ask whether humans use model-free versus model-based Baysian integration strategy (Figure 1).

### RSG task

Stimuli were presented on a fronto-parallel 23-inch display (distance: approximately 67 cm; refresh rate: 60 Hz; resolution: 1920 by 1200) and behavioral responses were registered using a standard Apple keyboard. Participants’ task was to measure a sample time interval (*t*_*s*_) between two visual flashes (‘Ready’ and ‘Set’) and subsequently initiate a delayed motor response (‘Go’) such that the produced interval relative to Set (*t*_*p*_) would match *t*_*s*_ as accurately as possible. Each trial began with the presentation of a central fixation point (FP; white circle, diameter: 0.5 degree in visual angle) along with a peripheral stimulus (white circle, diameter: 0.25 deg) that was presented 5 deg to the left of FP. We asked participants to maintain their gaze on FP throughout the trial but their eye positions were not monitored. The peripheral stimulus only served as a spatial reference and was otherwise irrelevant. After a random delay (exponentially distributed; mean of 500 ms with a minimum of 250 ms), Ready and Set flashes (white circle, diameter: 0.25 deg; flash duration: 50 ms) were presented in sequence to the right and top of FP. On each trial, *t*_*s*_ was sampled from the prior probability distribution (see below for details about the prior). After Set, participants had to produce a matching interval by pressing the spacebar on a keyboard, and received a graded numerical feedback (‘reward’) below FP based on the magnitude of the error (see below for details about the cost/reward function). Trials were separated by a fixed 500 ms inter-trial interval.

### Prior-cost metamers for RSG

On each trial, *t*_*s*_ was sampled from a prior distribution, and the numerical feedback was computed based on an experimentally imposed cost function. We measured performance in the context of two prior-cost pairs, the ‘original’ pair, which was introduced in the first session, and its ‘metamer’ that was additionally introduced on the remaining session.

We denote the original prior by *p*_*o*_(*t*_*s*_) and the original cost function by *C*_*o*_(*t*_*e*_,*t*_*s*_). For all participants, *p*_*o*_(*t*_*s*_) was a Gaussian distribution with a mean of 750 ms and a standard deviation of 144 ms (Figure 2c). *C*_*o*_(*t*_*e*_,*t*_*s*_) was a concave quadratic function of error (*t*_*e*_-*t*_*s*_) with a y-intercept of 100 such that participants received a maximum score of 100 when *t*_*e*_*=t*_*s*_ (Figure 2c). For errors larger than 1000 ms associated with negative scores, we truncated the cost function and set the score to 0. Mathematically, *p*_*o*_(*t*_*s*_) and *C*_*o*_(*t*_*e*_,*t*_*s*_) were defined as follows:

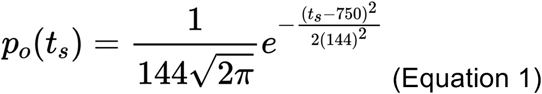

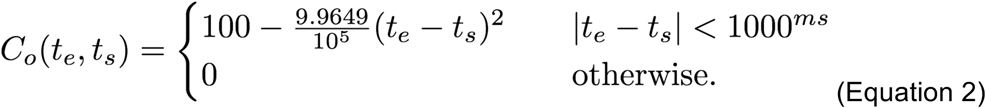

Following previous work^40^, we used a generative model to compute the ideal observer policy associated with the original prior-cost pair. We assumed that the ideal observer measurement of *t*_*s*_, denoted by *t*_*m*_, is perturbed by zero-mean Gaussian noise. Based on the scalar property of timing variability^75^, we additionally assumed that the standard deviation of noise scales with *t*_*s*_ with a constant of proportionality *w*_*m*_, which we refer to as the Weber fraction for measurement. Accordingly, we formulated the likelihood function, *p*(*t*_*m*_|*t*_*s*_) as *N*(*t*_*s*_,*w*_*m*_*t*_*s*_). We computed the ideal observer policy for the original pair, denoted by *f*_*o*_^*ideal*^, by integrating *p*(*t*_*m*_|*t*_*s*_) with *p*_*o*_(*t*_*s*_) and the original cost function by *C*_*o*_(*t*_*e*_,*t*_*s*_) and maximizing the expected reward:

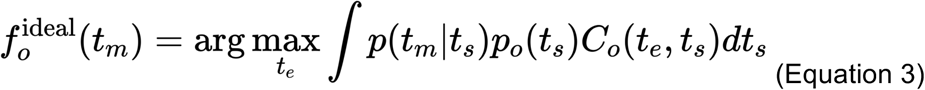

To compare the behavior of the participants to that of the ideal observer policy, we augmented the model to include production noise such that the produced interval, *t*_*p*_, was a sample from a Gaussian distribution with mean *t*_*e*_, and standard deviation *w*_*p*_*t*_*e*_ (*N*(*t*_*e*_,*w*_*p*_*t*_*e*_)) where *w*_*p*_ reflects additional scalar variability associated with the production^76^. We fit the ideal observer model to data from the first session to compare participants’ behavior to the optimal policy and to estimate each participant’s *w*_*m*_ and *w*_*p*_, which remained relatively stable throughout the entire experiment (Supplementary figure 8). Note that the scores participants received during the experiment were based on the error in *t*_*p*_, not *t*_*e*_, since *t*_*e*_ is not observable. However, using simulations, we verified that the expected cost function with *t*_*e*_ is similar to the cost function with *t*_*p*_ when the production noise is symmetric with respect to *t*_*e*_ (not shown). In computing the ideal observer policy, we did not take into account the fact that the quadratic cost function was truncated as errors were rarely within the truncated sidebands (mean±SD: 0.0064±0.0211% across subjects).

For the subsequent experimental sessions, we had to design a metamer for the original prior-cost pair. To do so, we considered a new prior, *p*_*m*_(*t*_*s*_) and a new cost function, *C*_*m*_(*t*_*e*_,*t*_*s*_) such that the ideal observer policy for the metamer, *f*_*m*_^*ideal*^ would be identical to *f*_*o*_^*ideal*^. Mathematically, this imposed the following equality constraint:

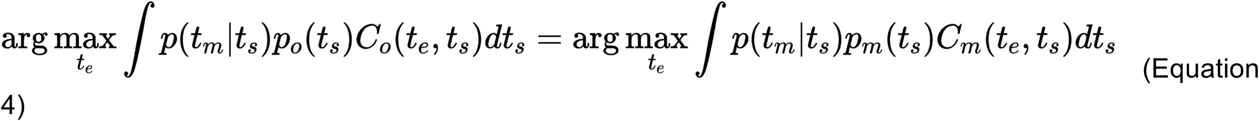

Evidently, there are an infinitude of solutions to this problem since we can freely adjust both *p*_*m*_(*t*_*s*_) and *C*_*m*_(*t*_*e*_,*t*_*s*_). For the purpose of our experiment, we wanted the original and its metamer to be generally similar but different enough to provide statistical power for detecting a transient learning effect. We focused on a specific form of manipulation that involved a shift in the cost function toward lower values, and commensurate adjustments to the prior to satisfy the constraint of generating a metamer. Accordingly, we defined *C*_*m*_(*t*_*e*_,*t*_*s*_) to be the same as *C*_*o*_(*t*_*e*_,*t*_*s*_) but shifted by an interval, *Δt*. Note that we favored this choice because of its computational simplicity but other manipulations to the cost function (e.g., introducing asymmetry) are also permissible.

Another factor we had to consider beyond matching the general form of the ideal observer policy was to make sure that the average score a participant received (the integral term in Equation 4) was similar between the original pair and its metamer. This was important because a sudden change in average reward (score) could signal a change in the environment and motivate learning a new policy. To address this point, we scaled *C*_*m*_(*t*_*e*_,*t*_*s*_) by a factor *k* whose value was adjusted so that the expected reward for the two pairs were the same. With these considerations in mind, the new cost function was formulated as follows:

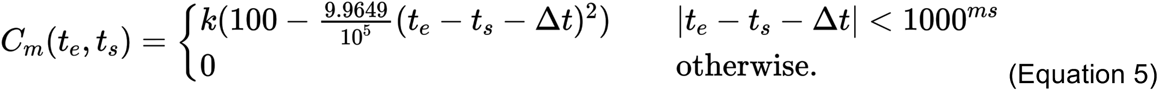

Using simulations (not shown), we determined that, depending on the Weber fraction, a shift in the cost function in the order of 100 to 200 ms (larger shifts for larger Weber fractions) was sufficient for detecting learning effects. Accordingly, we set *Δt* to 150 ms if the fitted *w*_*m*_ was less than 0.2, and 200 ms otherwise. Matching the average score also provided a participant-specific value for *k* (mean across participants: 0.73, SD: 0.06).

Having fixed the choice of *C*_*m*_(*t*_*e*_,*t*_*s*_), we next had to design *p*_*m*_(*t*_*s*_). To do so, we modeled *p*_*m*_(*t*_*s*_) as a Gaussian mixture model (GMM), Σ*w*_*k*_*N(µ*_*k*_, *σ*_*k*_), and estimated *w*_*k*_ and *σ*_*k*_ such that the new cost-prior pair was associated with the same ideal observer policy as the original pair (Equation 4; ‘fmincon’ function in MATLAB). To make this optimization process more tractable, we fixed the number of components in the GMM to 20 and fixed their corresponding mean to be equidistant between 350 to 1300 ms. Note that *p*_*m*_(*t*_*s*_) had to be customized for each participant separately because the exact shape *p*_*m*_(*t*_*s*_) depends on *w*_*m*_.

We then used the new prior-cost pair in the subsequent sessions. For the second and fourth sessions, we started the experiment with the original pair and then switched covertly to the metamer pair. For the third and fifth sessions, the order of the original and its metamer were switched. In all these sessions, the first prior-cost pair was present for the first 170 trials, and the other pair was present for the remaining trials (approximately 590 trials). More trials were dedicated to the post-transition so that we could observe the learning-related effects for longer periods.

### VMR task

Participants viewed the stimuli from above on a 21.5-inch Samsung SyncMaster SA200 display (distance: approximately 60 cm; refresh rate: 60 Hz; resolution: 1920 by 1080), and responded by controlling a custom manipulandum built from a stylus for a pen digitizer tablet (Wacom Intuous5 touch) with their hand occluded by the display. Participants’ task was to use a manipulandum to move a partially occluded cursor from its initial position at the center of the display to a target positioned over the circumference of a ring around the initial position. Each trial began with the presentation of a central point (FP; red square, size: 1 deg). To proceed, participants had to use the manipulandum to move the cursor, which at this stage of the trial was visible, to the center of the screen (over FP). As soon as the cursor entered an electronic circular window (diameter: 0.5 deg) around FP, a visual target (gray circle, contrast: 40%, diameter: 1 deg) was presented briefly (duration: 500 ms) on a circular ring (radius: 10 deg, thickness: 0.1 deg, contrast: 50%) around FP. The ring remained visible throughout the trial. The position of the target was chosen from a uniform distribution between 80 to 100 deg with its midpoint aligned to 12 o’clock over the circle. Afterwards, participants had to move the cursor to the remembered location of the flashed target. The cursor was made invisible as soon as it passed the electronic window, and remained invisible throughout its movement except for a brief reappearance (gray circle, contrast: 40%, diameter: 0.5 deg, duration: 100 ms) when it was 3 deg from FP. Participants were asked to move the cursor smoothly with no interruption.

Critically, while the cursor was invisible and before its reappearance at 3 deg from FP, we applied an angular rotation (*x*_*s*_) to the cursor relative to the hand position. On each trial, *x*_*s*_ was taken from a prior distribution (see below for details about the prior). Participants had to use their knowledge about the prior distribution and the visual feedback along the path to infer *x*_*s*_, and maneuver the cursor to produce a counter rotation (*x*_*p*_) that matches *x*_*s*_ as accurately as possible. We defined *x*_*p*_ as the angular distance between the target and hand position over the ring. Similar to RSG, participants received a graded numerical feedback below FP based on the magnitude of the error (see below for details about the cost/reward function). Trials were separated by a fixed 500 ms inter-trial interval. For VMR, we asked participants to use their dominant hand throughout the experiment.

### Prior-cost metamers for VMR

The original prior, *p*_*o*_(*x*_*s*_) and cost function *C*_*o*_(*x*_*e*_,*x*_*s*_) were governed by the following equations:

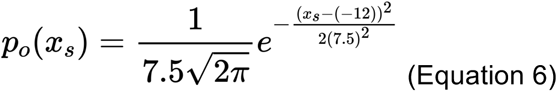

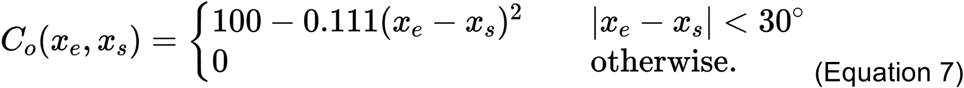

To compute the ideal observer policy for the VMR task, we assumed that the measured angle, *x*_*m*_ is subject to zero-mean Gaussian noise, and formulated the likelihood function, *p*(*x*_*m*_|*x*_*s*_) as *N*(*x*_*s*_,*σ*_*m*_). In this setting with a Gaussian prior and a Gaussian likelihood function, *f*_*o*_^*ideal*^ is a simple linear mapping between *x*_*e*_ and *x*_*m*_ as follows:

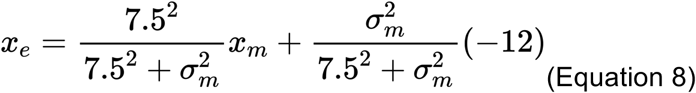

To compare the participants behavior to the ideal observer, we additionally assumed that the produced angular correction, *x*_*p*_, was also subject to zero-mean Gaussian noise with standard deviation, *σ*_*p*_; i.e., *x*_*p*_∼*N*(*x*_*e*_,*σ*_*p*_). We fit the ideal observer model to data from the first session to compare participants’ behavior to the optimal policy and to estimate each participant’s *σ*_*m*_ and *σ*_*p*_, which remained relatively stable throughout the entire experiment (Supplementary figure 9). Note that we defined the cost function in terms of error in *x*_*e*_, and computed the scores based on the error in *x*_*p*_. Similar to the RSG task, we verified that the expected cost function with *x*_*e*_ is similar to the cost function with *x*_*p*_ when the production noise is symmetric with respect to *x*_*e*_ (not shown).

Our procedure for devising a metamer for the original prior-cost pair was as follows: we formulated *p*_*m*_(*x*_*s*_) as a Gaussian distribution that was shifted by 10 deg, and *C*_*m*_(*x*_*e*_,*x*_*s*_) as a quadratic function that was shifted in the opposite direction, as follows:

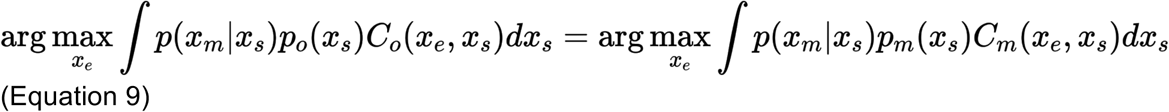

Note that the design of the metamer for the VMR is significantly easier than the RSG task because, in the VMR task, changing the mean of the prior only alters the intercept of the linear policy and not its slope, which can be readily compensated by an opposite shift in the cost function. We computed the requisite shift in the cost function (*Δx*) analytically by matching *f*_*o*_^*ideal*^ to *f*_*m*_^*ideal*^ as follows:

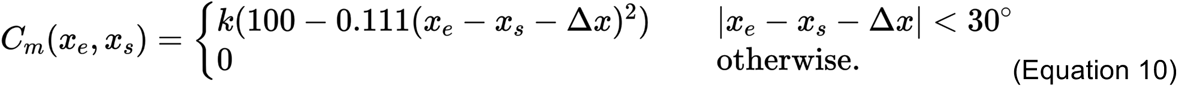

where *Δx* can be computed as follows:

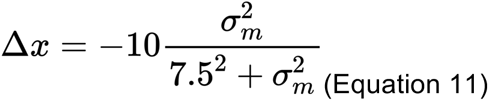

Finally, we used the original and metamer prior-cost pairs in the subsequent sessions using the same procedure as in the RSG task.

### Models and Analysis

All analyses were performed using custom code in MATLAB (Mathworks, Inc.). We first removed outlier trials for each data set across all participants and sessions. We applied different algorithms to the two tasks as their response profile was inherently different (i.e., nonlinear metronomic function for the RSG task and linear policy for the VMR). For the RSG task, we excluded trials in which the relative error, defined as (*t*_*p*_-*t*_*s*_)/*t*_*s*_, deviated more than 3 standard deviations from its mean (mean: 0.50%; SD: 0.28% across subjects). For the VMR task, we first fitted a linear regression model relating *x*_*p*_ and *x*_*s*_, and excluded trials for which the error from the linear fit was more than 3.5 times larger than the median absolute deviation (mean: 3.4%; SD: 2% across data sets). We verified that outlier trials were not concentrated immediately after the switch between the prior-cost pairs.

#### Decision policy for the model-free (H_1_) and model-based (H_2_) optimal strategies in the RSG task

For the original prior-cost pair, which uses a quadratic cost function, the optimal policy is to choose the mean of the posterior:

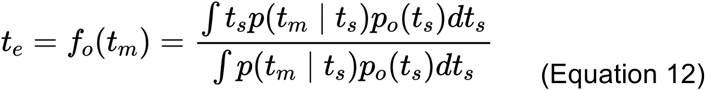

This function captures the asymptotic optimal policy for both the model-free and model-based strategies because the two pairs were designed to be metamers. The same optimal policy can also be written in terms of the *p*_*m*_*(t*_*s*_) as follows:

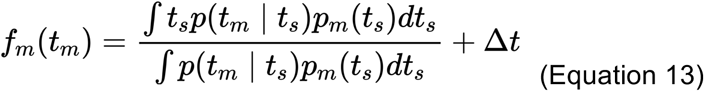

Note that the shift of the cost function, *Δt*, adds a constant offset to the policy, which is compensated by the change in the prior.

#### Decision policy for prior-sensitive (H_3_) and cost-sensitive (H_4_) suboptimal strategies

Under H_3_, the observer only learns the new prior and assumes that there has been no change in the quadratic cost function. The corresponding decision policy is therefore to use the mean of the posterior under the new prior:

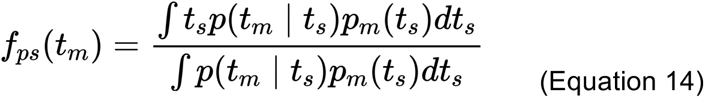

Under H_4_, the observer only learns the new cost function and assumes that there has been no change in the Gaussian prior. The corresponding decision policy can be computed as follows:

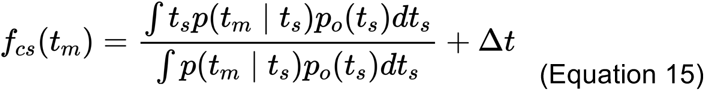

The policy for the prior-sensitive and cost-sensitive models has the same form as *f*_*m*_(*t*_*m*_) and *f*_*o*_(*t*_*m*_), respectively, but their intercepts are different from the optimal policies (Figure 2d).

#### Model comparison after switch

To distinguish between H_1_, H_2_, H_3_, and H_4_, we compared the log-likelihood of the data under the four models (Equations 12-15) and asked which model better captures behavior of the participants immediately after the switch between the two prior-cost pairs. The likelihood for each trial (superscript *i*) under each model (*j*) can be computed as follows:

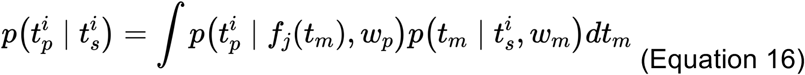

The first and second term within the integral correspond to the noisy generative process of production and measurement, respectively (*w*_*p*_ and *w*_*m*_ for its Weber faction). *f*_*j*_*(t*_*m*_) can be replaced by *f*_*o*_ (H_1_/H_2_: model-free and model-based optimal policy), *f*_*ps*_ (H_3_: prior-sensitive), and *f*_*cs*_ (H_4_: cost-sensitive) to test the data under the corresponding hypotheses.

#### Decision policies and model comparisons in the VMR task

Equations 12-16 also apply to the VMR task by substituting *t*_*s*_, *t*_*m*_, *t*_*e*_, *t*_*p*_ with *x*_*s*_, *x*_*m*_, *x*_*e*_, *x*_*p*_, respectively.

#### Fitting subjective prior and cost function for re-learning strategy

To better characterize the relearning process after the prior-cost switch, we fitted the behavior after switch using an ideal observer model in which the prior and cost function could change dynamically across trials. This exercise allowed us to examine the time course of relearning and provided a basis for later analyses where we compared the learning rates between the prior and cost function. To do so, we assumed participants used a Gaussian prior and a quadratic cost function, and treated the mean of the prior, *µ*, and shift in the cost function, *Δt*, as free parameters of the model (in addition to the variance of the prior and the two Weber fractions for measurement and production; Supplementary Figure 4) that were fitted based on the behavioral data after the switch. The two Weber fractions as additional parameters did not change the result in Figure 4 as a simpler model without free *w*_*m*_ and *w*_*p*_ led to qualitatively similar results (not shown). We estimated the model parameters by maximizing the likelihood of data *t*_*p*_ and *t*_*s*_ (Equation 16 with the decision policy *f*_*j*_*(t*_*m*_) from Equation 13; ‘fminsearch’ function in MATLAB) under the model. We used a running window of 100 trials, which led to 1 pre-transition and 4 post-transition estimates of *µ* and *Δt*. We did not fit a subjective model to the VMR data as we could not reject the model-free strategy (Figure 7a).

Finally, we compared different models (model-free, model-based, prior-sensitive, cost-sensitive) across all data sets using Bayesian Information Criteria (BIC=-2×*L*+*N*_*param*_×*log*(*N*_*obs*_) where *L* is the likelihood of data given model parameters, *N*_*param*_ is the number of free parameters in the model, *N*_*obs*_ is the number of data points, i.e., trials). For model-free, prior-sensitive, and cost-sensitive models, we used the first session’s *w*_*m*_ and *w*_*p*_ estimates, which remained relatively stable across sessions (Supplementary figure 8). Therefore, *N*_*param*_ was zero for all models except the subjective re-learning model (5 per each data set, 100-trial window).

#### Analysis of learning rates and simulation

The subjective re-learning model allowed us to capture how the internal prior and cost function is updated after the transition. We fitted an exponential function to the time course of the subjective *µ* and *Δt* to quantify and compare the corresponding learning rates for each participant. To make the fits more robust, we constrained the initial and final values of *µ* and *Δt* to match the experimentally imposed values for the pre- and post-transition (*µ*_*pre*_, *Δt*_*pre*_ and *µ*_*post*_, *Δt*_*post*_, respectively). With this consideration, we only needed to fit the parameter associated with the learning rate (*τ*_*µ*_, *τ*_*Δt*_; Figure 5a), as follows.

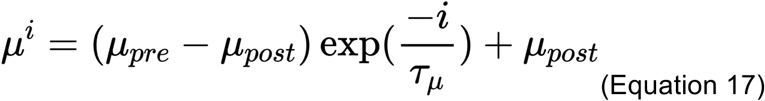

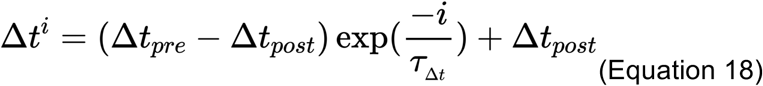

Where *µ*^*i*^, *Δt* ^*i*^ are the prior mean and cost shift in trial *i* after the transition. We estimated *τ*_*µ*_, *τ*_*Δt*_ by maximizing the likelihood of data *t*_*p*_ and *t*_*s*_ (Equation 16 with the decision policy *f*_*j*_*(t*_*m*_) from Equation 13; ‘fminsearch’ function in MATLAB). Similar to the subjective model fitting, we allowed both the mean and variance of the prior to change throughout learning.

We also performed simulations to examine how the learning rates for *µ* and *Δt* influence performance (Figure 5f). The procedure for each simulation was as follows: 1) we chose specific values for the learning rates, *τ*_*µ*_ and *τ*_*Δt*_; 2) we computed *µ* and *Δt* as a function of time governed by those learning rates; 3) we computed the moment by moment value of expected reward (*EC*) as a function of the instantaneous values of *µ* and *Δt* as follows:

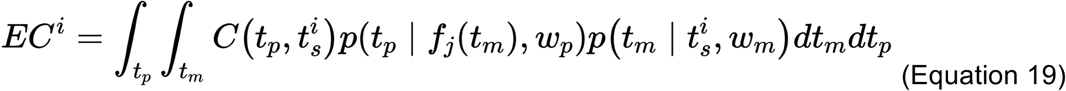

In this equation, we computed *EC*^*i*^ (expected cost in trial *i*) in the context of the metamer prior-cost pair for an observer with *w*_*m*_ =0.14 and *w*_*p*_ = 0.08, close to averages across subjects (Supplementary Figure 8). In this equation, different values of *µ* and *Δt* influence *EC* through the decision policy *f*_*j*_*(t*_*m*_) (Equation 13). We also measured the overall average reward (marginalizing over time) for different choices of *τ*_*µ*_ and *τ*_*Δt*_.

## Supporting information

HS_MJ_priorCost_Supplementary

## Acknowledgments

H.S. is supported by the Center for Sensorimotor Neural Engineering. M.J. is supported by NIH (NINDS-NS078127), the Sloan Foundation, the Klingenstein Foundation, the Simons Foundation, the McKnight Foundation, the Center for Sensorimotor Neural Engineering, and the McGovern Institute.

## Author contributions

H.S. and M.J. conceived the experiments. H.S. collected and analyzed the data. M.J. supervised the project. H.S. and M.J. wrote the manuscript.

## Data availability

Data collected and used in this study will be made publicly available at https://jazlab.org/resources/.

## Code availability

Code used in this study will be made publicly available at https://jazlab.org/resources/.

