## Supplementary material for "Metamers of Bayesian computation": HS_MJ_priorCost_Supplementary

**Supplementary Figures**

**Figure S1.** Comparison of ideal observer model to behavior in RSG task.

**Figure S2.** Data from all participants in RSG task.

**Figure S3.** Test of transient suboptimal policy under pre-transition prior.

**Figure S4.** Time course of parameters in subjective prior-cost model.

**Figure S5.** Comparison of observer models and behavior for task performance in RSG.

**Figure S6.** Data from all participants in VMR task.

**Figure S7.** Comparison of observer models and behavior for task performance in VMR.

**Figure S8.** Time course of Weber fractions in RSG.

**Figure S9.** Time course of measurement and production noise in VMR.

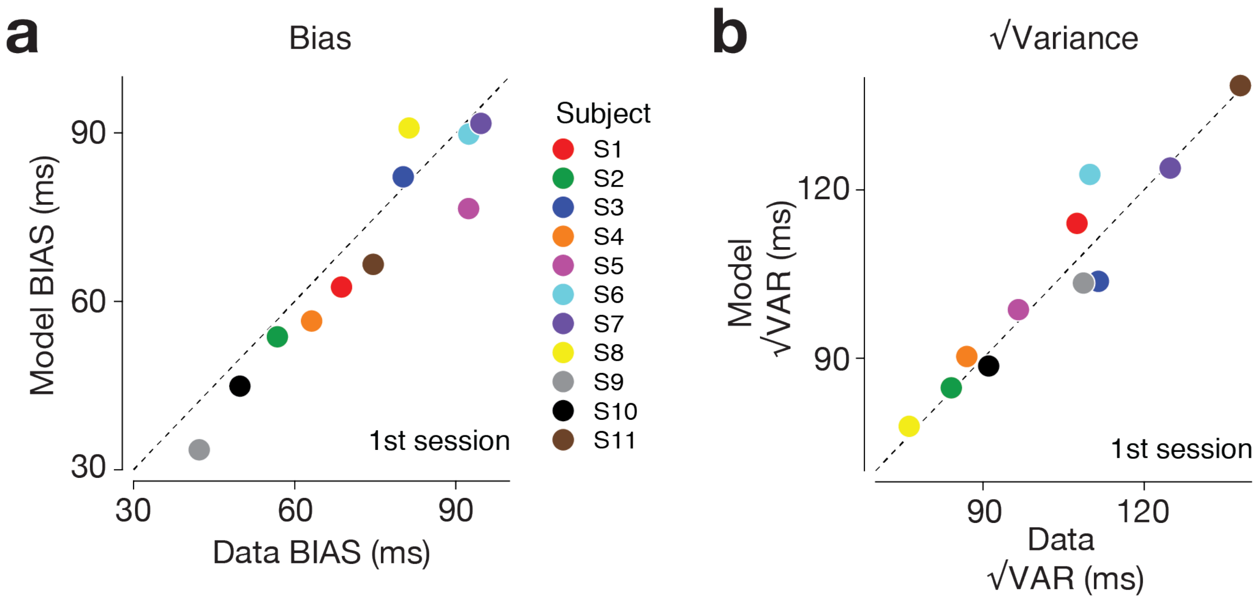


**Figure S1.** Comparison of ideal observer model to behavior in RSG task in the first session. We compared participants' behavior in RSG task to the prediction of the ideal observer model using two criteria, BIAS (**a**) and √VAR (**b**). These can be formally defined as follows:

$$\mathrm{BIAS} = \sqrt{E\left[bias^2\right]} = \sqrt{\frac{1}{N}\sum_{i=1}^{N} bias_i^2}$$

$$\sqrt{VAR} = \sqrt{E[var]} = \sqrt{\frac{1}{N}\sum_{i=1}^{N} var_i}$$

$N$ is the number of bins that we used to group the continuously sampled $t_s$ (approximately 14). We computed $bias_i$ and $var_i$ (bias and variance of the $i$-th bin) as follows:

$$bias_i = E_i[t_p] - t_s^i$$

$$var_i = E[t_p - E_i[t_p]]$$

where $E_i[t_p]$ is average $t_p$ for a given bin and $t_s^i$ is the average $t_s$ within that bin. We also measured the same statistics for the ideal observer associated with each participant. To do so, we simulated the generative model to perform the same exact experiment (same $t_s$ values) using $w_m$ and $w_p$ fits to each participant. Participants' behavior was consistent with that of the ideal observer model.

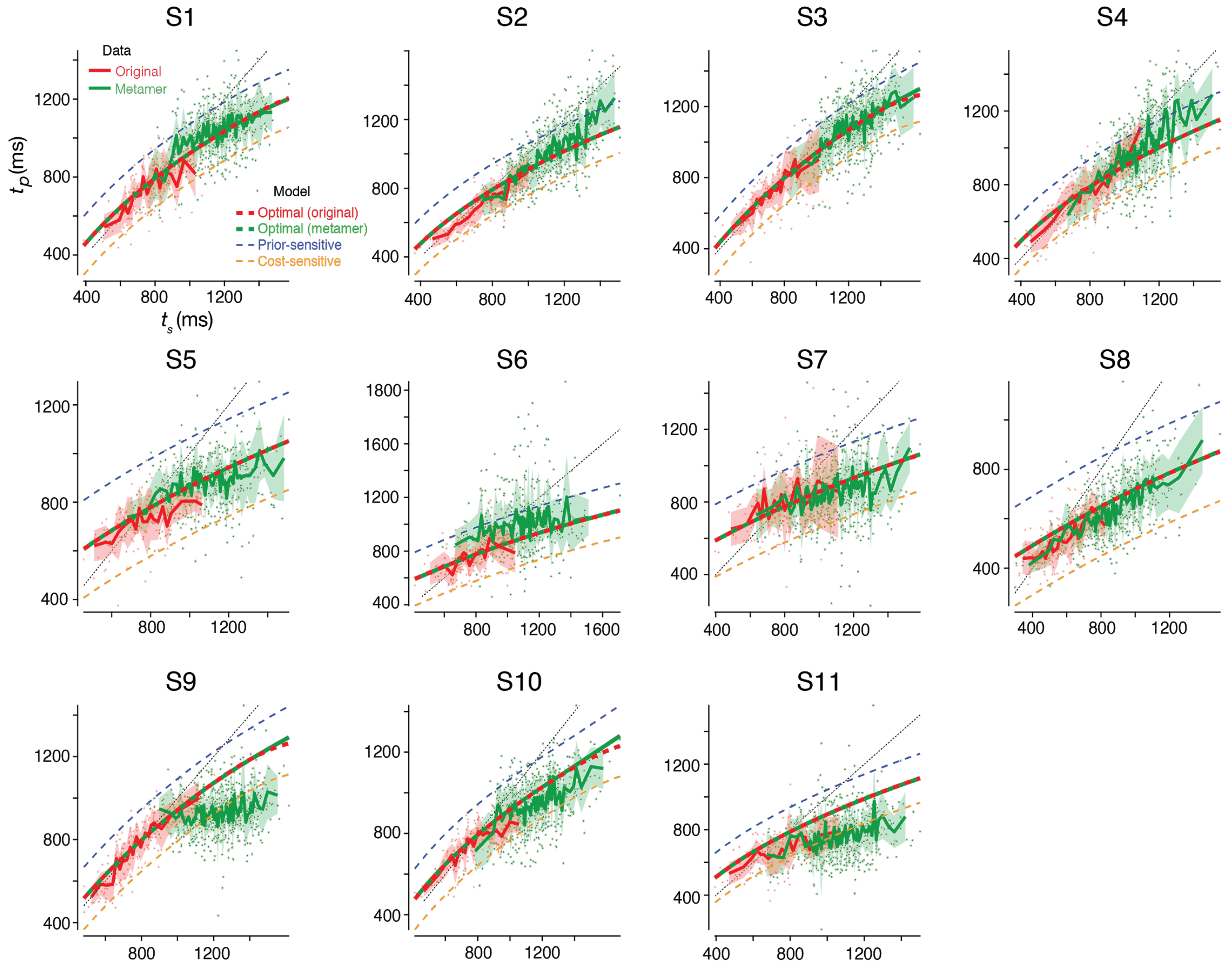


**Figure S2.** Data from all participants in the RSG task. Results are shown in the same format as in Figure 2d.

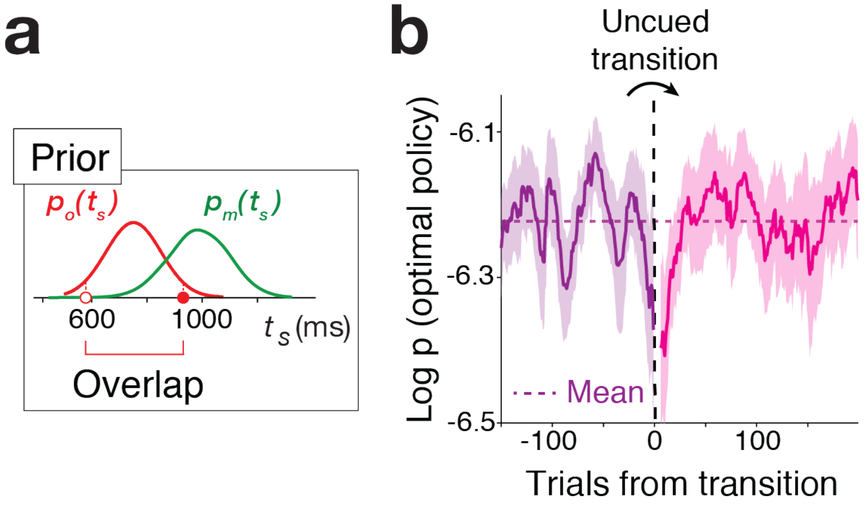


**Figure S3.** The transient deviation from optimal policy for the subset of stimuli that overlap between the original and metamer priors. One potential reason behind the transient deviation from optimal policy in Figure 3a is that some of the $t_s$ values in the mentamer condition were not present in the original condition. To address this concern, we reanalyzed the probability of the optimal policy for the subset of trials whose $t_s$ values overlapped with the range of $t_s$ values before the transition. (**a**) For the transition from the original to the metamer (second and fourth sessions), we analyzed trials in which $t_s$ was between the minimum (empty red circle) and maximum $t_s$ (filled red circle) tested in the first session. For the transition from the metamer to the original (third and fifth sessions), we analyzed trials in which $t_s$ was between the minimum and maximum $t_s$ tested in the metamer condition of the second session. (**b**) Same as in Figure 3a for the overlapping values of $t_s$. Results indicate that participants' decision policy deviated from the optimality even for previously experienced $t_s$ values validating our original conclusion from Figure 3a.

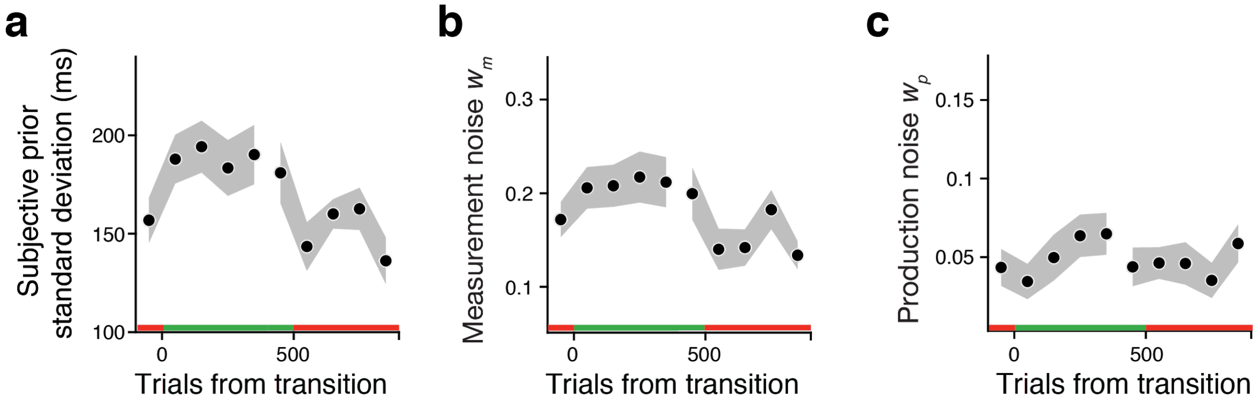


**Figure S4.** Time course of parameters in subjective prior-cost model. The (**a**) standard deviation of the subjective prior, (**b**), measurement Weber fraction ($w_m$), and (**c**) production Weber fraction ($w_p$) were relatively stable during transitions between the original and metamer conditions. Results are shown in the same format as Figure 4a (mean of the prior) and 4b (shift of the cost function). The original prior has SD of 144ms and the SD of the metamer (Gaussian mixture distribution) was 163.85 ± 19.25 (mean ± SD across participants).

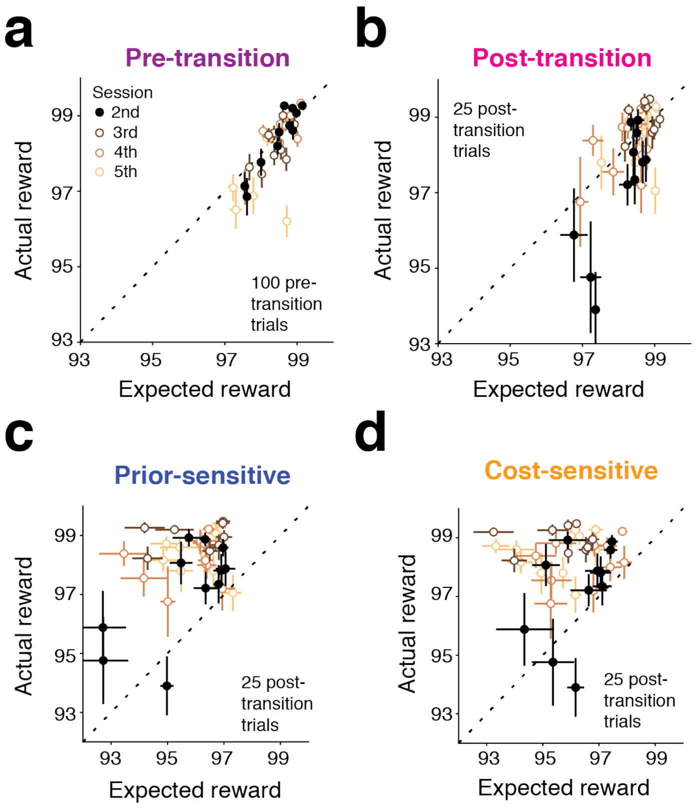


**Figure S5.** Comparison of observer models and participants for task performance in RSG. We used task performance (i.e., reward score) as another metric to compare participants' behavior to different models (optimal, prior-sensitive, and cost-sensitive). We compared the actual reward each participant received to the expected reward (Equation 19 in Method) under each model (Equation 12-15 in Method) using the Weber fractions derived from fits of the ideal observer to that participants' behavior in the 1st session. (**a**) Before the transition, actual reward was similar to the expected reward under the ideal observer model (circles: individual participants and sessions, error bar: SEM across 100 trials). (**b**) After the transition, the actual reward remained near the optimal level ($p=0.14$, one-tailed sign-rank test), although participants performed significantly worse than the ideal observer for the 2nd session ($p=0.012$, one-tailed sign-rank test; black circles), in which they experienced the transition for the first time. We chose 25 post-transition trials for this analysis based on the time course of p(optimal policy) in Figure 3a. (**c**,**d**) Subjects showed better performance than expected from either the prior-sensitive model ($p<0.001$, one-tailed sign-rank test) and the cost-sensitive model ($p<0.001$, one-tailed sign-rank test), consistent with findings in Figure 3b,c.

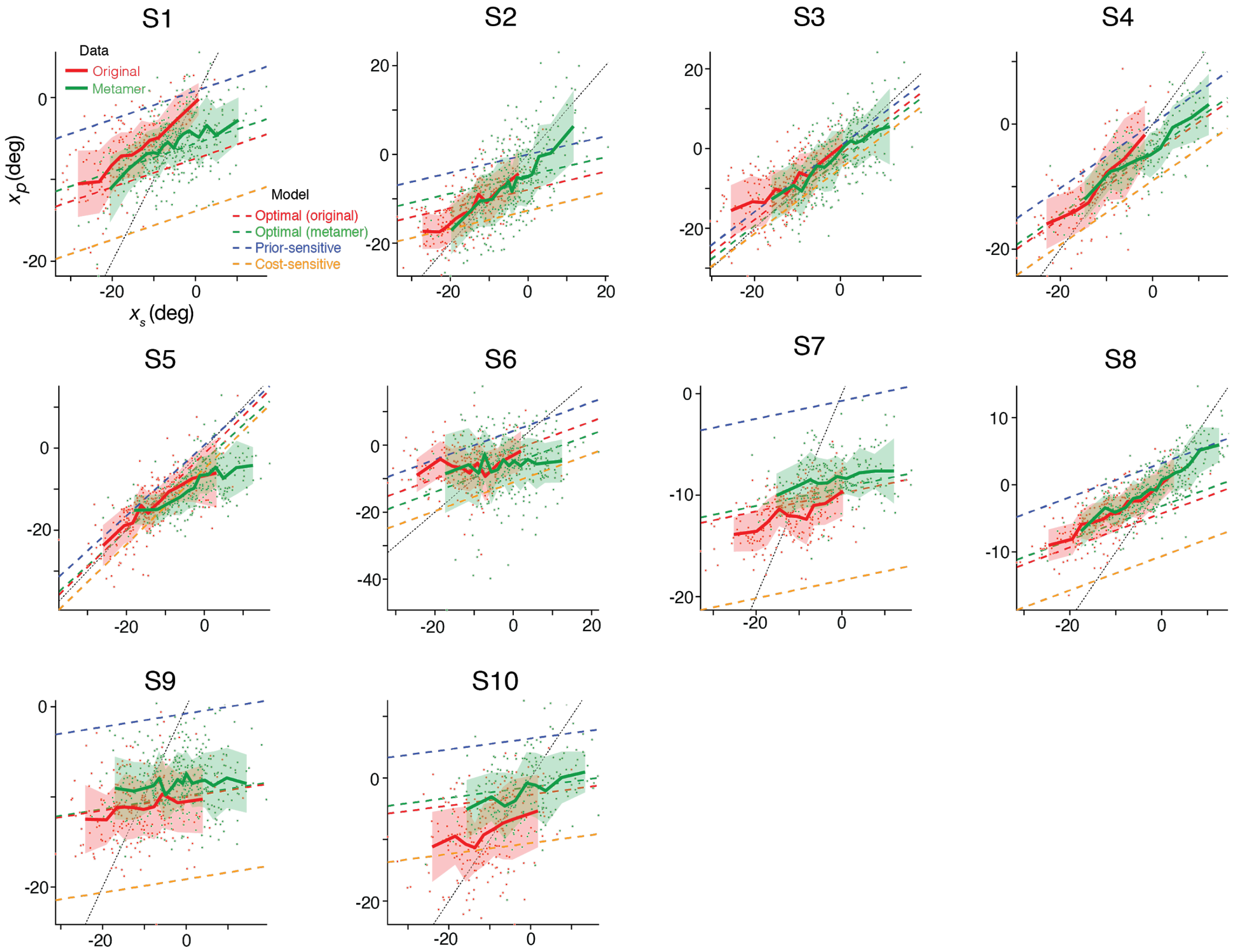


**Figure S6.** Data from all participants in the VMR task. Results are shown in the same format as in Figure 6d.

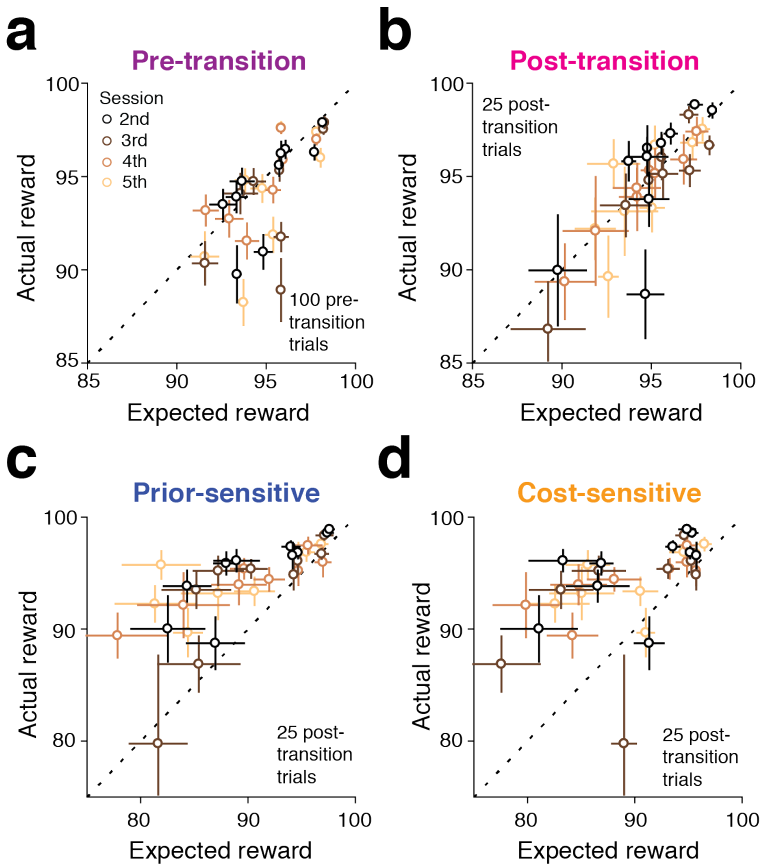


**Figure S7.** Comparison of observer models and participants for task performance in VMR in the same format as in Figure S5. We compared the actual reward each participant received to the expected reward (Equation 19 in Method) under each model (Equation 12-15 in Method) using the measurement and production noise parameters derived from the fits of the ideal observer to that participants' behavior in the 1st session. (**a**) Before the transition, actual reward was similar to the expected reward under the ideal observer model. (**b**) After the transition, task performance overall remained near the optimal level (p=0.061, one-tailed sign-rank test). Subjects showed better performance than expected from either the prior-sensitive model (p<0.001, one-tailed sign-rank test) and the cost-sensitive model (p<0.001, one-tailed sign-rank test), consistent with findings in Figure 7b,c.

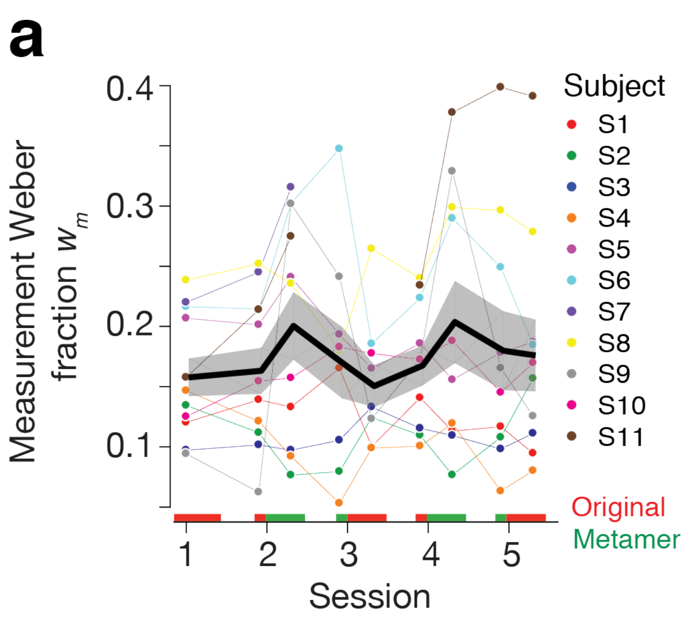


**Figure S8.** Time course of Weber fractions in RSG. Weber fractions were fitted to pre- and post-transition epochs (shown over the abscissa) separately.

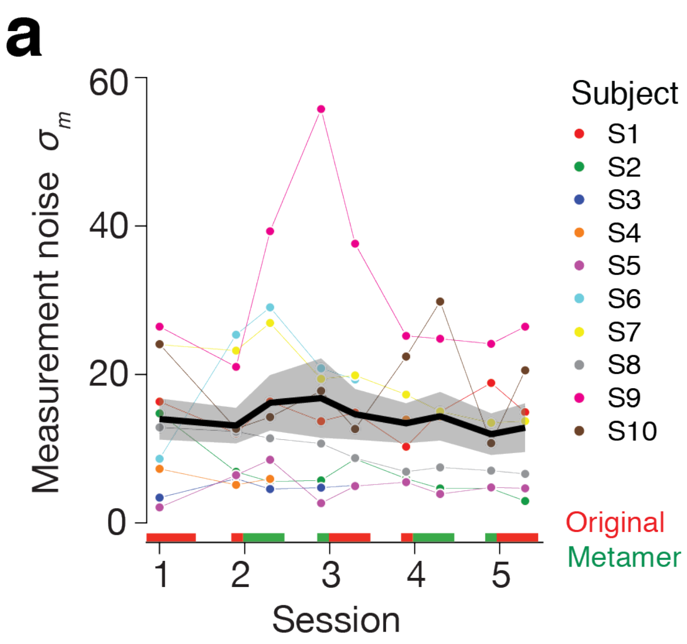


**Figure S9.** Time course of measurement and production noises in VMR. $\sigma_m$ and $\sigma_p$ were fitted to pre- and post-transition epochs (shown over the abscissa) separately.
